# Comparison of gene-by-gene and genome-wide short nucleotide sequence based approaches to define the global population structure of *Streptococcus pneumoniae*

**DOI:** 10.1101/2024.05.29.596230

**Authors:** Alannah C. King, Narender Kumar, Kate C. Mellor, Paulina A. Hawkins, Lesley McGee, Nicholas J. Croucher, Stephen D. Bentley, John A. Lees, Stephanie W. Lo

**Author notes:** Corresponding Author Contact: Alannah C. King, Corresponding Author Contact: Stephanie W. Lo. co-first author who contributed equally (Alannah C. King, Narender Kumar). co-senior author who contributed equally (John A. Lees, Stephanie W. Lo).

## Abstract

Defining the population structure of a pathogen is a key part of epidemiology, as genomically related isolates are likely to share key clinical features such as antimicrobial resistance profiles and invasiveness. Multiple different methods are currently used to cluster together closely-related genomes, potentially leading to inconsistency between studies. Here, we use a global dataset of 26,306 *S. pneumoniae* genomes to compare four clustering methods: gene-by-gene seven-locus multi-locus sequencing typing (MLST), core genome MLST (cgMLST)- based hierarchical clustering (HierCC) assignments, Life Identification Number (LIN) barcoding, and k-mer-based PopPUNK clustering (known as GPSCs in this species). We compare the clustering results with phylogenetic and pan-genome analyses to assess their relationship with genome diversity and evolution, as we would expect a good clustering method to form a single monophyletic cluster that has high within-cluster similarity of genomic content. We show that the four methods are generally able to accurately reflect the population structure based on these metrics, and that the methods were broadly consistent with each other. We investigated further to study the discrepancies in clusters. The greatest concordance was seen between LIN barcoding and HierCC (Adjusted Mutual Information Score = 0.950), which was expected given that both methods utilise cgMLST, but have different methods for defining an individual cluster and different core genome schema. However, the existence of differences between the two methods show that the selection of a core genome schema can introduce inconsistencies between studies. GPSC and HierCC assignments were also highly concordant (AMI = 0.946), showing that k-mer based methods which use the whole genome and do not require the careful selection of a core genome schema are just as effective at representing the population structure. Additionally, where there were differences in clustering between these methods, this could be explained by differences in the accessory genome that were not identified in cgMLST. We conclude that for *S. pneumoniae*, standardised and stable nomenclature is important as the number of genomes available expands. Furthermore, the research community should transition away from seven-locus MLST, and cgMLST, GPSC, and LIN assignments should be used more widely. However, to allow for easy comparison between studies and to make previous literature relevant, the reporting of multiple clustering names should be standardised within research.

**Data summary:** Genome sequences are deposited in the European Nucleotide Archive (ENA); accession numbers. Metadata of the pneumococcal isolates in this study have been submitted as a supplementary file and are also available on the Monocle Database available at https://data.monocle.sanger.ac.uk/. The authors confirm all supporting data, code and protocols have been provided within the article or through supplementary data files.

**Impact Statement:** Using a global dataset of *S. pneumoniae* genomes allows us to thoroughly observe and analyse discrepancies between different clustering methods. Whilst all methods in this study are used to cluster *S. pneumoniae* genomes, no study has yet thoroughly compared the clustering results and discrepancies. This work summarises the strengths and weaknesses of the different methods and highlights the need for consistency between studies.

## Introduction

*Streptococcus pneumoniae* (or pneumococcus) is a clinically important human pathogen responsible for a range of infectious diseases, including otitis media, pneumonia, and meningitis. It was estimated to have caused 829,000 (95% uncertainty intervals 682,000– 1,010,000) deaths worldwide in 2019 [1]. The polysaccharide capsule surrounding the cell of the pneumococcus is the basis of a traditional typing scheme which has been used to separate isolates into groups (i.e. serotypes) [2]. Although the pneumococcal capsule is the major virulence factor and vaccine target, the disease potential [3], transmissibility [4], antimicrobial resistance [3], and responses to vaccine-induced protection [5] are also related to the genetic composition of the strain [3]. Therefore, defining the population structure of pneumococci lays the foundation for epidemiological investigation [4,6] and assessing the impact of clinical interventions such as vaccine implementation and antimicrobial use for prophylaxis or treatment of disease [7].

Due to high levels of recombination, defining the population structure of pneumococci is particularly challenging. Since 1998, multi-locus sequence typing (MLST) has been widely used to capture the genetic variation of seven housekeeping genes (*aro*E, *gdh, gki*, *rec*P, *spi*, *xpt* and *ddl*) to classify the isolates into sequence types (STs) [8]. For each locus, a unique sequence is assigned an arbitrary and unique allele number. The designations for the seven loci are incorporated into an allelic profile (e.g. 1-1-1-1-1-1-1), and the ST (e.g.ST1) is assigned based on the allelic profile. STs can further be grouped into clonal complexes (CCs) based on the goeBURST algorithm [9]. The cutoff between CCs is usually defined as variation at one locus by most users [9]. The CCs are then named after the founder ST identified by the algorithm. For example, the founder ST of CC156 is ST156.

MLST is portable and designed to accommodate the conflicting signals of vertical and horizontal genetic transfer to infer evolutionary relationships [10]. Additionally, it is straightforward to compute, and does not require whole-genome sequencing. Over the years, limitations of the MLST scheme have been observed as an increasing number of pneumococcal genomes become available. Firstly, it is known that the absence or disruption of genes within the MLST scheme can prevent ST assignment, and preliminary data has identified this in *gki* and *xpt* [11]. Secondly, high recombination frequency in *gki*, *gdh*, *recP*, *spi* and *ddl* can obscure vertical genetic transfer, resulting in over-clustering due to convergent MLST profiles in phylogenetically disparate isolates [12,13]. For example, *ddl* has been seen in linkage with *pbp2b*, which is a recombination hotspot [14]. In addition, our previous study has identified that larger collections of genomes are more vulnerable to spurious connections when assigning CCs [3]. Similar observations have also been reported in other bacterial species [15]. Thirdly, CC assignments can vary between collections, making the nomenclature inconsistent between studies. Finally, MLST has limited resolution capabilities - when applied to closely related isolates belonging to the same ST, the limited number of loci involved can lead to under-clustering as it is unable to identify fine-grained evolutionary events that distinguish strains within a recent epidemic [16,17].

As more genomes became available, another gene-by-gene approach, core-genome MLST (cgMLST), was devised to expand the number of genes included in the MLST analysis. Here, analysis of the gene content of a selected subset of genomes is used to determine a set of genes that reflect a core genome. Clustering into groups is then based upon the allelic profile of these core genes [18], often upwards of 1,000 compared to the seven housekeeping genes in a MLST scheme, therefore increasing the resolution significantly and making better use of the bacterial genome compared to seven allele MLST. The allelic profiles of the core genes are most commonly used for clustering, but they can also be used to construct phylogenetic trees using the SplitsTree4 package [19] or hierarchical clustering (HierCC) [20]. However, these phylogenies do not account for the nucleotide diversity within each locus as the allelic profile from the cgMLST scheme does not contain this information [18]. Although publicly available core genomes schemes exist for many bacteria, cgMLST is not widely used in *S. pneumoniae* [18,20], which could be in part due to how laborious defining a core genome schema is and the difficulties in having it be universally adopted. Using keywords “Streptococcus pneumoniae cgMLST”, only three publications were found (searched conducted on 10^th^ Apr 2024) [21–23]. There is also whole genome MLST (wgMLST), which creates an allelic profile for each genome based on every open reading frame, instead of only seven loci (MLST) or the core genome (cgMLST) only. However this was not investigated in this study as research in *S. enterica* has shown clusters assigned by wgMLST to be highly similar to those from cgMLST assignments [24,25].

Recently, a related typing system – Life Identification Numbers, or LIN – has been proposed which makes use of cgMLST to provide each *S. pneumoniae* genome with a barcode based on the allelic similarity between it and other genomes in the dataset [18]. This approach has been used to study *Klebsiella*, which similarly to *S. pneumoniae* has high genomic diversity [26]. One benefit of the barcoding system is that each lineage has defined clusters within it, providing a high level of resolution beyond the lineage level. Additionally, it automatically keeps names from legacy schemes [26]. However, as it is based on cgMLST, it still does not account for nucleotide diversity within each locus, and requires the creation of a scheme of core genes [18]. Different core gene sets and different cut-offs for allelic similarity may lead to different clustering results, and thus inconsistent nomenclature.

An alternative reference-free approach based on k-mer similarity, as opposed to allelic profiles, has been proposed. Here, short nucleotide sequences of length k, known as k-mers, are used to measure core and accessory genome similarity and cluster them. For example, Population Partitioning Using Nucleotide K-mers (PopPUNK) [27] - has been used to define genomic definitions of pneumococcal lineage (or Global Pneumococcal Sequence Cluster, GPSC) on a global collection of ∼20,000 genomes [3]. This approach grouped isolates that had shared evolutionary history into a GPSC based on variations across the entire genome, in a scalable fashion with standard nomenclature. Generally, GPSCs had high concordance with ST/CC, and GPSC assignments often encompassed related CCs [3,28].

With the growing number of pneumococcal genomes available in the Global Pneumococcal Sequencing (GPS) project, discrepancies between GPSC and ST/CC have increasingly been observed. In this study, we compared the clustering results from the gene-by-gene based approaches (MLST, cgMLST and LIN barcoding), and a k-mer based approach (PopPUNK/GPSC). We then conducted an in-depth investigation into the discrepancies using phylogenetic and pan-genome analyses. Given that clustering should reflect the population structure, we would expect a successful clustering method to produce clusters that are monophyletic in the species-wide tree, showing within-cluster similarity of genomic content.

## Methods

### Genome Collection and *In Silico* Serotyping

In the GPS project, a global collection of 26,306 *S. pneumoniae* genomes were sequenced from 57 countries, 1989-2020 [29]. These published genomes passed quality control as previously described [6]. Briefly, the sequence reads had a base quality >20, *S. pneumoniae* was the predominant species in Kraken with >60% species-specific reads, number of contigs assembled (<500), total assembled length (between 1.9 Mb and 2.3 Mb), mapping coverage for *S. pneumoniae* reference (ATCC 700669, Accession number: FM211187 [30]) >60%, overall coverage depth >20X and the number of heterozygous single nucleotide polymorphism (SNP) sites <220. *In silico* serotyping was carried out using SeroBA with default parameters [31].

### Assignment of ST and CC

The pneumococcal genomes in this study were assigned STs using MLSTcheck version 2.1.17 [7]. The allelic profile of seven housekeeping genes (*aro*E, *gdh, gki*, *rec*P, *spi*, *xpt,* and *ddl*) were input for clustering using goeBURST algorithm based on a cutoff of single locus variants. The goeBURST algorithm is implemented in Phyloviz v2.0 [8]. The CC was named as per the ST identified as the primary founder by goeBURST.

### GPSC

For each genome in this study, GPSC were assigned using PopPUNK v2.6.0 [4] with the reference database GPS_v6. The reference database contains 42,163 pneumococcal genomes and 1,112 GPSCs.

### cgMLST and HierCC

The genome assemblies were used to create cgMLST using chewBBACA v3.2.0 [32]. A subset of 100 randomly chosen genomes were first used to create the schema and then the remaining 26,206 genome assemblies were used to detect the presence of alleles defined in the schema [32]. Together, cgMLST profile for the entire collection was created which was composed of 1,248 genes. The cgMLST profiles were then used to perform further clustering using HierCC v1.27 [33] with an allelic distance of 620, as this provided the highest silhouette score with HCCeval [33].

### LIN Barcodes

LIN barcodes were accessed on the 2^nd^ of Feb 2023 [18] and incorporated into the dataset via ERR numbers. 765 genomes in the LIN database did not have ERR numbers. Samples within our dataset that did not have a LIN barcode were not used for LIN-related analysis, but our full dataset of 26,306 genomes was used for CC, HierCC, and GPSC analysis. To compare to lineage assignments, the first three numbers of LIN barcodes were used and reported to define lineage level [18].

### Single Cluster Phylogenetic and Pan-genome Analyses

For the selected sets of genomes within a single cluster (either one GPSC, one CC, or one HierCC), we used panaroo v1.3.3 [12] to identify core and accessory genome content. In tandem, Gubbins v3.2.1 [34] [35] was used to build a recombination free phylogeny for each single cluster of genomes analysed (e.g. a single GPSC, or a single CC). The reference sequence used for each cluster was that of the dominant GPSC. This tree along with core and accessory genome content was visualised in Phandango [36].

### Species-Wide Phylogenetic Analyses

Pseudo-genomes were created using samtools mpileup after mapping the genomes to the *S. pneumoniae* reference genome (ATCC 700669, Accession number: FM211187). SNP sites were identified from the pseudo-genomes using snp-sites version 2.5.1 [37]. FastTree v2.1.10 [38] was used to construct a maximum likelihood phylogenetic tree using the SNP sites across all 26,306 genomes. FastTree [38] used a generalised time reversible model and gamma rate substitution. The species-wide tree and associated data can be visualised on Microreact (https://microreact.org/project/clusteringcomparison).

### Statistical analysis

The Adjusted Mutual Information (AMI) score measures the agreement between clustering methods after adjusting for the agreement expected between random clustering. The score ranges between -1 to 1, whereby -1 indicates complete disagreement, 0 that the clustering is no better than random, and 1 that there is complete agreement between clustering methods. The function AMI from the R package aricode was used [39].

The mean (μ) and standard deviation (σ) of the number of clusters produced by method A within a single cluster of method B were calculated in R.

## Results

### Genome collection summary

The global collection of pneumococcal genomes (n=26,306) was derived from isolates causing invasive disease (n=12,063), non-invasive disease (n=630) and asymptomatic colonisation (n=7,948) from sixty countries, 1989-2020 (supplementary table). Of 26,306 pneumococcal genomes, 81 *in silico* serotypes were identified. A subset of 17,361 genomes within our database had associated LIN barcodes from the Pneumococcal Genome Library, and so these were incorporated.

### Overview of Clustering Methods

For 26,306 pneumococcal genomes, 4,766 STs were identified and grouped into 1,656 CCs. Of these, 801 (48%) CCs were only composed of a single ST (i.e. were singletons). 25,812 cgMLSTs were identified in the same set of pneumococcal genomes, which were clustered into 606 HierCCs in which 146 (24%) were singletons. Using PopPUNK, 830 GPSCs were identified of which 272 (30%) were singletons (**Table 1**). Given the differences in the number of total clusters, the mean number of sequences within each cluster varied accordingly; methods that led to fewer total clusters ended up with more genomes per cluster (**Table 1**, **Figure 1A**). This decreases the resolution of the method and potentially leads to difficulties in tracking small-scale outbreaks of disease. On the other hand, as the number of total clusters increases, so does the proportion of singletons, and so the benefits that come from clustering are reduced.

**Figure 1.**
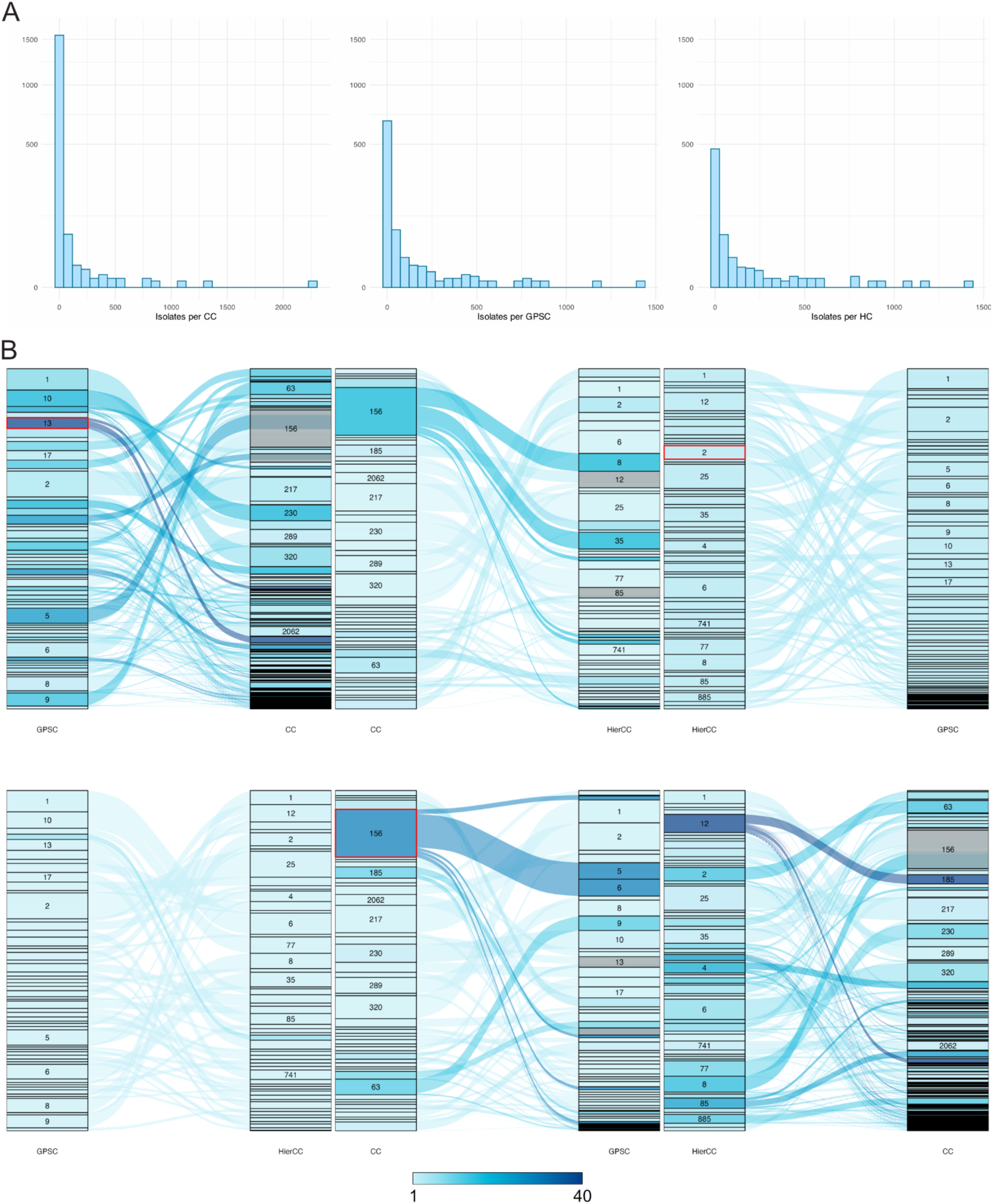
Number of genomes in one cluster for each method. (A) Many CCs contain just one or very few genomes, compared to GPSC and HierCC clusters. (B) Sankey plots to visualise the difference between methods. Each left bar represents clusters with over 100 genomes determined by one method, shown being distributed into clusters defined with a second method to the bar on the right. The colour shows the number of clusters produced by method two for a single cluster from method one; the darker the blue, the more clusters it has been split into. Clusters analysed in further depth in the results are highlighted in red.

**Table 1.** Statistical analysis of the results of the three different clustering methods. Mean (μ) and standard deviations (σ) for the number of cluster type B within calculated using R.

|  |  | CC | HierCC | GPSC |
| --- | --- | --- | --- | --- |
| *Total Clusters* |  | 1,656 | 606 | 830 |
| *No. of singletons (%)* |  | 801 (48%) | 146 (24%) | 272 (30%) |
| *Max cluster size* |  | 2,265 | 1,417 | 1,417 |
| *Min cluster size* |  | 1 | 1 | 1 |
| *Members per cluster* | $\mu$ | 15.9 | 43.4 | 31.7 |
| | $\sigma$ | 86.65 | 135.84 | 109.19 |
| *A single CC* | $\mu$ | | 1.04 | 1.07 |
| | $\sigma$ | | 0.33 | 0.70 |
| *A single HierCC* | $\mu$ | 2.83 | | 1.38 |
| | $\sigma$ | 4.77 | | 1.43 |
| A single GPSC | $\mu$ | 2.13 | 1.01 | |
| | $\sigma$ | 2.95 | 0.08 | |

AMI scores were used to compare the level of agreement between different clustering methods. The AMI score between CC and GPSC assignments was 0.869, showing high concordance between the two methods; 97% (1,608/1,656) of CCs were composed of genomes belonging to a single GPSC. After removing singleton CCs (n=801), this changed to 94% (807/855). The AMI score between HierCC and GPSC clustering was 0.946, showing that there is greater concordance between HierCC and GPSC assignments than CC and GPSC assignments. In the subset of the data which had assigned LIN barcodes, the AMI score between HierCC and LIN Lineages was 0.950, showing almost complete concordance but still a degree of variability likely introduced through the selection of the core genome schema and different definitions of lineage. Interestingly, this shows that despite both using core genome schemes, HierCC and LIN lineages are as similar to each other as HierCC and GPSC assignments are to each other. Overall, CC assignments were the most different from the other methods (**Figure 1B**).

None of the methods produced identical results to each other. To investigate these differences further, we selected examples for further analysis where a single cluster produced using one method was split into multiple different clusters by another method. For example, there were 48 (3%) CCs containing genomes belonging to multiple (>1) GPSCs and 285 (35%) GPSCs with genomes belonging to >1 CCs.

### CCs containing Multiple GPSCs

To learn more about CCs containing multiple GPSCs, we investigated all the CCs containing at least 500 genomes. There were eight CCs that met this criterion (CC156, n = 2,265; CC217, n = 1,299; CC320, n = 1,085; CC230, n = 864; CC289, n = 754; CC63, n = 742; CC185, n = 537; CC2062, n = 520). Of these, four consisted of a single GPSC (CC217, CC320, CC289, CC2062) and so were not investigated further. The other four CCs contained multiple different GPSCs (CC156, n = 23; CC63, n = 10; CC185, n = 9; CC230, n = 2;). As CC156 contained the greatest number, we investigate this example in further detail here.

CC156 contained 23 different GPSCs, with GPSC5 and GPSC6 being the dominant GPSCs. This corresponded to nine HierCCs, and 14 LIN lineages (**Figure 2A**). Within these LIN lineages, there were seven different superlineages, suggesting large and substantial differences between the genomes (**Figure 2A**). The genomes within CC156 are a polyphyletic group, and so do not cluster in the species-wide phylogeny and are spread across multiple distant branches of the tree, supporting the assignment to different superlineages and lineages. When coloured by GPSC, it is possible to see that individual clades within the tree have been assigned to a different GPSC (**Figure 2B**), and similar is true for HierCC and LIN lineages (**Figure 2C and 2D**).

**Figure 2.**
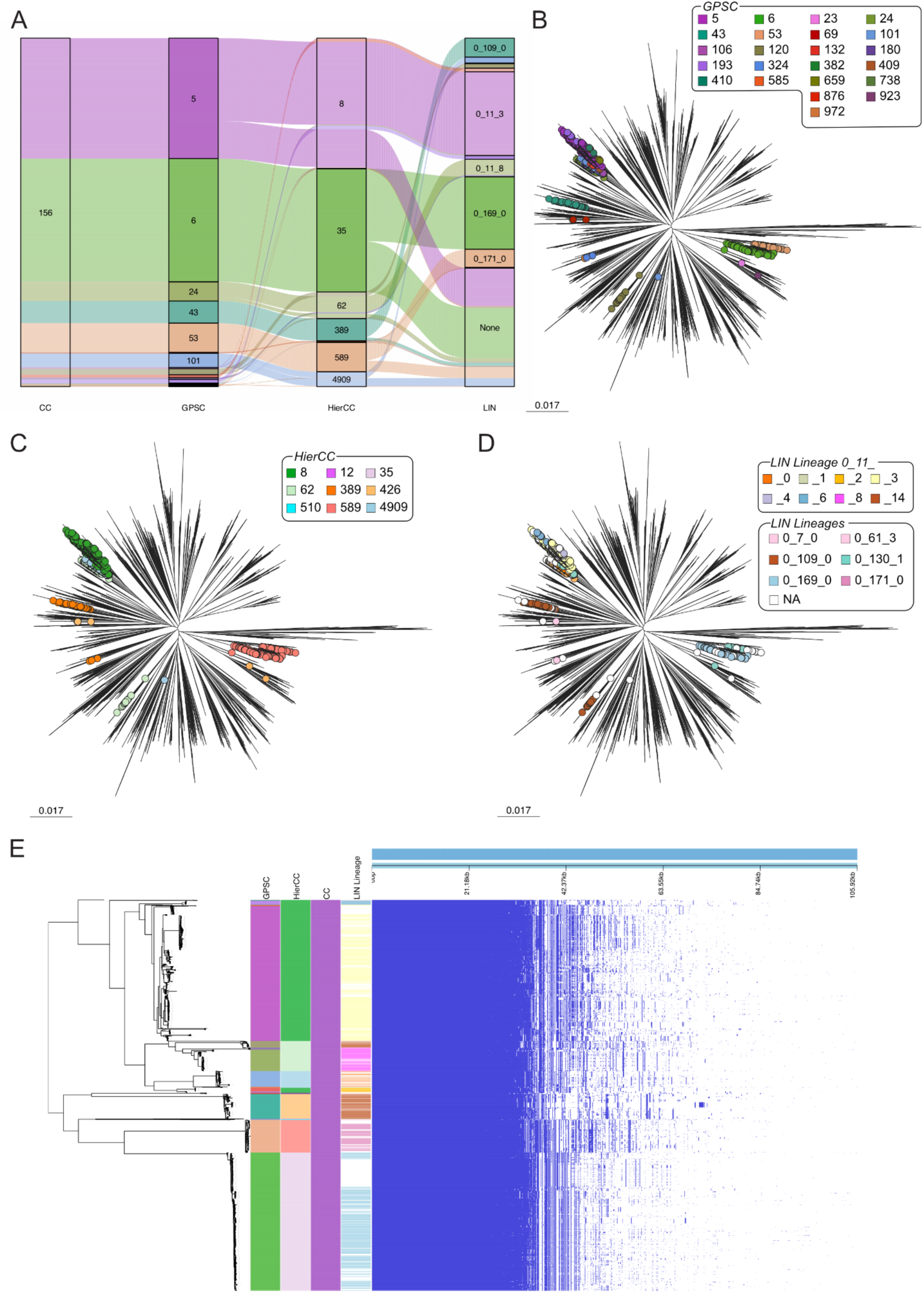
Analysis of genomes assigned to CC156. (A) Within CC156, there were multiple CCs, HierCCs, and LIN lineages. (B) In a species-wide tree, the genomes within CC156 were spread across the whole phylogeny. GPSCs (C) and LIN lineages (D) were better able to cluster according to the phylogeny. (E) Pan-genome analysis of CC156 with a recombination free tree. There are clear differences in genomic content between genomes, broadly correlating with GPSC, HierCC, and LIN Lineage assignments.

Pan-genome analysis of CC156 shows the concordance between GPSC, HierCC, and LIN lineage assignments of the genomes within CC156. Visual differences in the gene content can be seen between the different major HierCCs, GPSCs, and LIN lineages, supporting even further the subdivision of CC156 into multiple groupings. However, for the assignments of minor groupings, it is difficult to conclusively determine if this subdivision is biologically relevant and valid.

Together, this shows that the few genes used to define a CC are likely to group together distantly related genomes, creating difficulties when trying to track epidemics and when discussing lineages. In this case, CC156 contains genomes within ST156, which has been identified as a genotype of high concern due to its rapid spread and high resistance to penicillin and cotrimoxazole [30].

### GPSCs Containing Multiple CCs

We investigated the GPSCs containing at least 500 genomes to identify GPSCs containing multiple CCs. There were nine GPSCs that contained over 500 genomes, and each of these contained multiple CC (**Table 2**),

**Table 2.** GPSCs with over 500 genomes with them, and the number of CCs contained within each.

| GPSC | Number of Genomes | Number of CCs |
| --- | --- | --- |
| 1 | 1,166 | 9 |
| 2 | 1,417 | 5 |
| 5 | 835 | 24 |
| 6 | 803 | 4 |
| 8 | 765 | 3 |
| 9 | 713 | 16 |
| 10 | 897 | 19 |
| 13 | 592 | 35 |
| 17 | 524 | 4 |

GPSC13 is equivalent to HierCC 85, however it is split into 35 CCs (**Figure 3A**). As this is the greatest number of CCs seen within a single GPSC in this dataset, we performed a thorough analysis of it. The genomes within GPSC13 formed a single monophyletic clade in the species-wide phylogeny, reflecting a shared ancestry that would be expected from a single GPSC (**Figure 3B**). The subtree of GPSC13 showed that the two dominant CCs, CC473 and CC2285, formed distinct groupings (**Figure 3C**). However, there were many minor CCs scattered within two, with 68.6% of the CCs only containing a single genome. 88.6% of the CCs within GPSC13 were singletons, containing a single ST that was not found elsewhere in the database. GPSC13 contained six different LIN Lineages, all of which were within the 0_21 super lineage (**Figure 3D**), reflecting their close relatedness. These lineages sat in clusters in the subtree.

**Figure 3.**
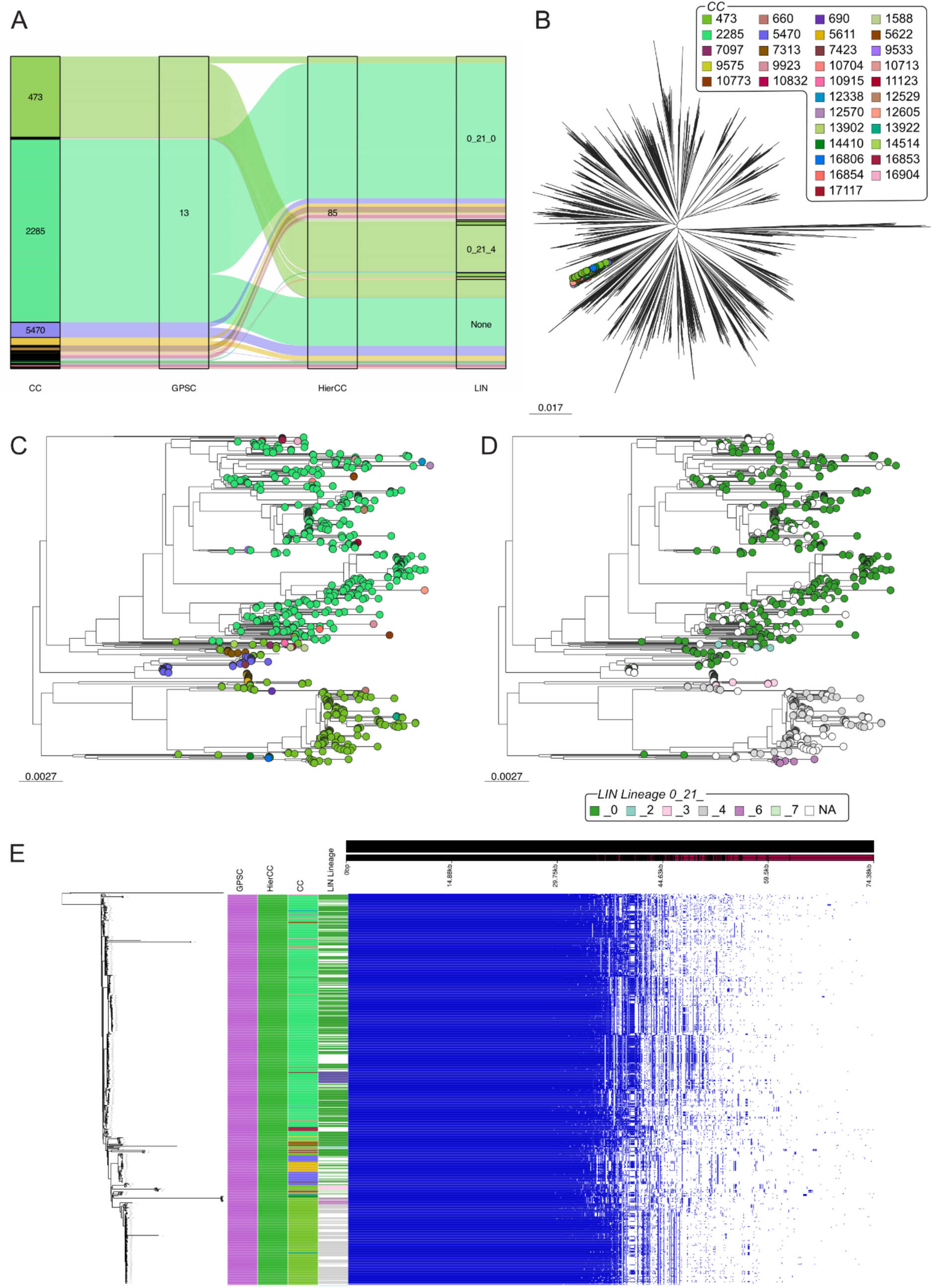
Analysis of genomes assigned to GPSC13. (A) Within the single GPSC and HC, there were multiple CCs and LIN lineages. (B) In a species-wide tree, the genomes within GPSC13 clustered together. There was limited clustering at the CC (C) level and LIN lineage (D) level. (E) Pan-genome analysis with a recombination-free tree of GPSC13 genomes showed high similarity between the genomes, not supporting the high number of CC assignments.

The same phylogenetic clustering was seen in the recombination-free GPSC13 phylogeny (**Figure 3E**). Pan-genome analysis of the genomes within GPSC13 showed them to be similar to each other, with a core genome (>95%) size of 1743 genes. The different CCs and LIN lineages do not correspond to any clear visual changes in genome content from the pangenome analysis (**Figure 3E**), suggesting that the further subdivision is not reflective of significant changes in the genome.

We performed the same analysis on the other GPSCs containing multiple CCs, and the figures can be found in the supplementary. Interestingly, we continued to see the same pattern of GPSCs and HierCCs being concordant with each other, whilst there are multiple CCs present. Often, the genomes are dominated by one CC with multiple minor CCs containing very few genomes. This suggests that CC assignments are likely to be noisy.

### HierCCs Containing Multiple GPSCs

83.3% (505/606) of HierCCs contained a single GPSC, and therefore 16.6% (101/606) HierCCs contained multiple GPSCs. We identified the five HierCCs that contained the most GPSCs for further analysis (HierCC2, n = 791; HierCC12, n = 1063; HierCC8, n = 933; HierCC390, n = 257; HierCC885, n = 515). The greatest number of GPSCs was seen in HierCC2, which contained 17 GPSCs and 27 CCs (**Figure 4A**), and so we analysed this further.

**Figure 4.**
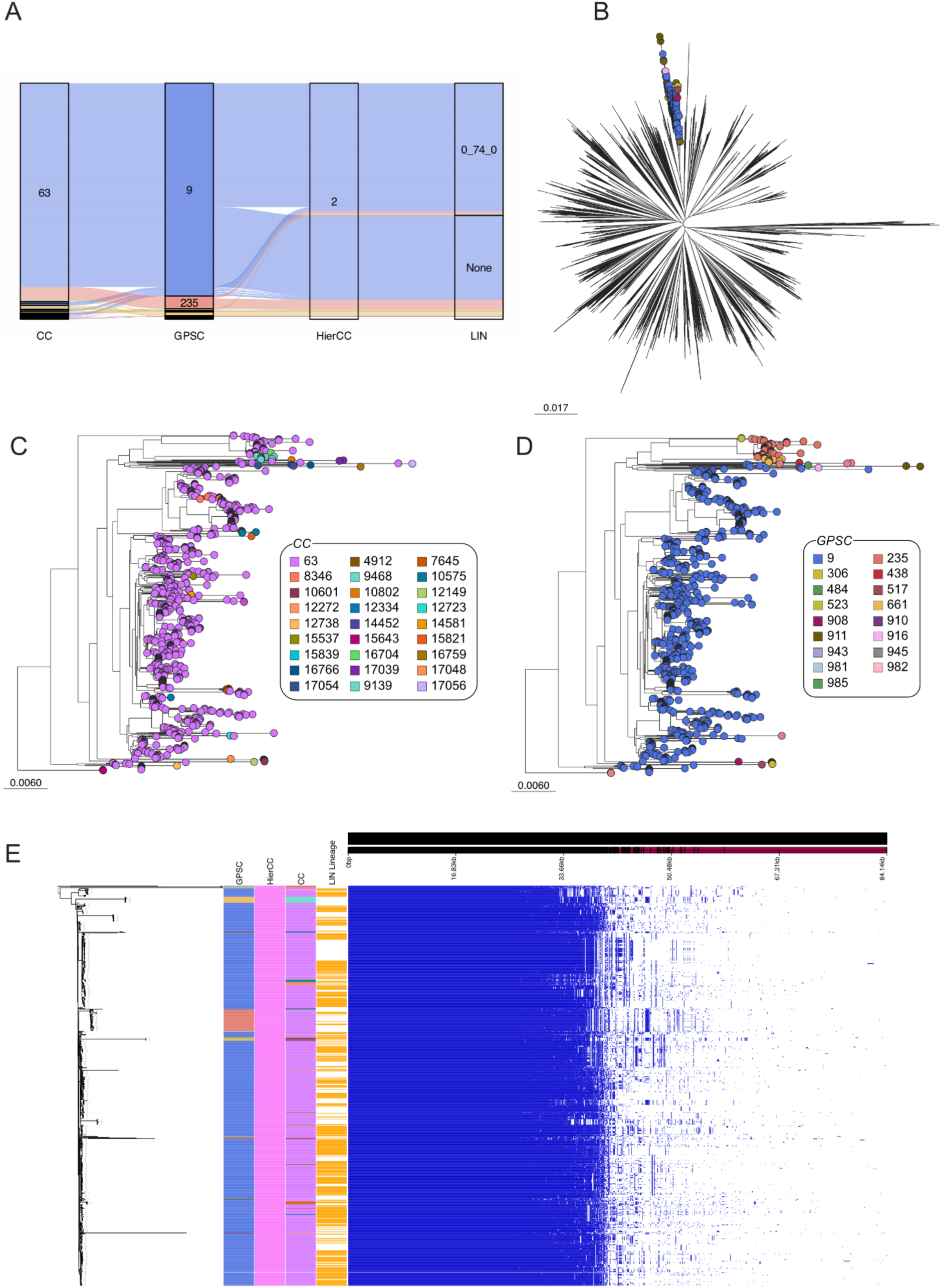
Analysis of genomes assigned to HierCC2. (A) Within the one HierCC, there were multiple GPSCs and CCs. (B) In a species-wide tree, the genomes within HierCC2 were in the same branch. (C) CC63 is the dominant CC, with many minor CCs spread throughout. (D) The two main GPSCs formed two clusters, with other minor GPSCs. (E) Pan-genome analysis with a recombination-free tree of HierCC2 genomes showed high similarity between the genomes. Differences in the accessory genome rationalise the existence of GPSC235 as a separate cluster.

The genomes in HierCC2 only belonged to a single LIN lineage, 0_74_0, showing the similarity between the two cgMLST methods despite the schema differences. All the genomes within HierCC2 sat within a single branch of the species-wide phylogeny (**Figure 4B**). Colouring the subtree by CC showed that the vast majority of genomes were assigned to CC63, with many other minor CCs spread throughout (**Figure 4C**). 66.7% (18/27) of CCs within HierCC2 were represented by only a single genome. These single genomes being assigned to different CCs, despite clustering with other CC63 isolates throughout the phylogeny, indicates that solely using the CC designation may lead to clonal expansions or outbreaks being missed. Colouring the subtree by GPSC showed that the majority of samples belonged to GPSC9 or GPSC235, with each of these forming distinct clusters (**Figure 4D**). Genomes belonging to other GPSCs can be seen, often clustering together.

The recombination-free phylogeny of HierCC2 genomes also showed clustering by GPSCs (**Figure 4E**). Pan-genome analysis supported the assignment of the genomes within GPSC235 into their own cluster, as shown by clear differences in the accessory genome compared to other genomes (**Figure 4E**). These genes include some from transposons, recombinases, and hypothetical proteins amongst others (**Supplementary Table 2)**. This difference was not identified by the HierCC, CC or LIN lineage assignments, reflecting whether or not they account for the accessory genome.

We performed the same analysis on the other HierCCs that contain multiple GPSCs, and the figures can be found in the supplementary (**Supplementary figures 11, 12, 13, and 14**).

### GPSCs Containing Multiple HierCCs

There were very few GPSCs containing multiple HierCCs; the mean number of HierCCs per GPSC was 1.01. Only six GPSCs contained two HierCCs (GPSC12, GPSC45, GPSC112, GPSC272, GPSC326, GPSC511). Each of these GPSCs sat within the same branch of the phylogeny, showing high relatedness (**Figure 5**). Individual subtrees showed clustering of the two HierCCs into different branches, suggesting that HierCCs provide slightly higher resolution clustering than GPSCs (**Figure 5**). Given the few genomes within each GPSC, pan-genome analysis was not performed as reasonable conclusions cannot be drawn from so few samples about the validity of the clustering.

**Figure 5.**
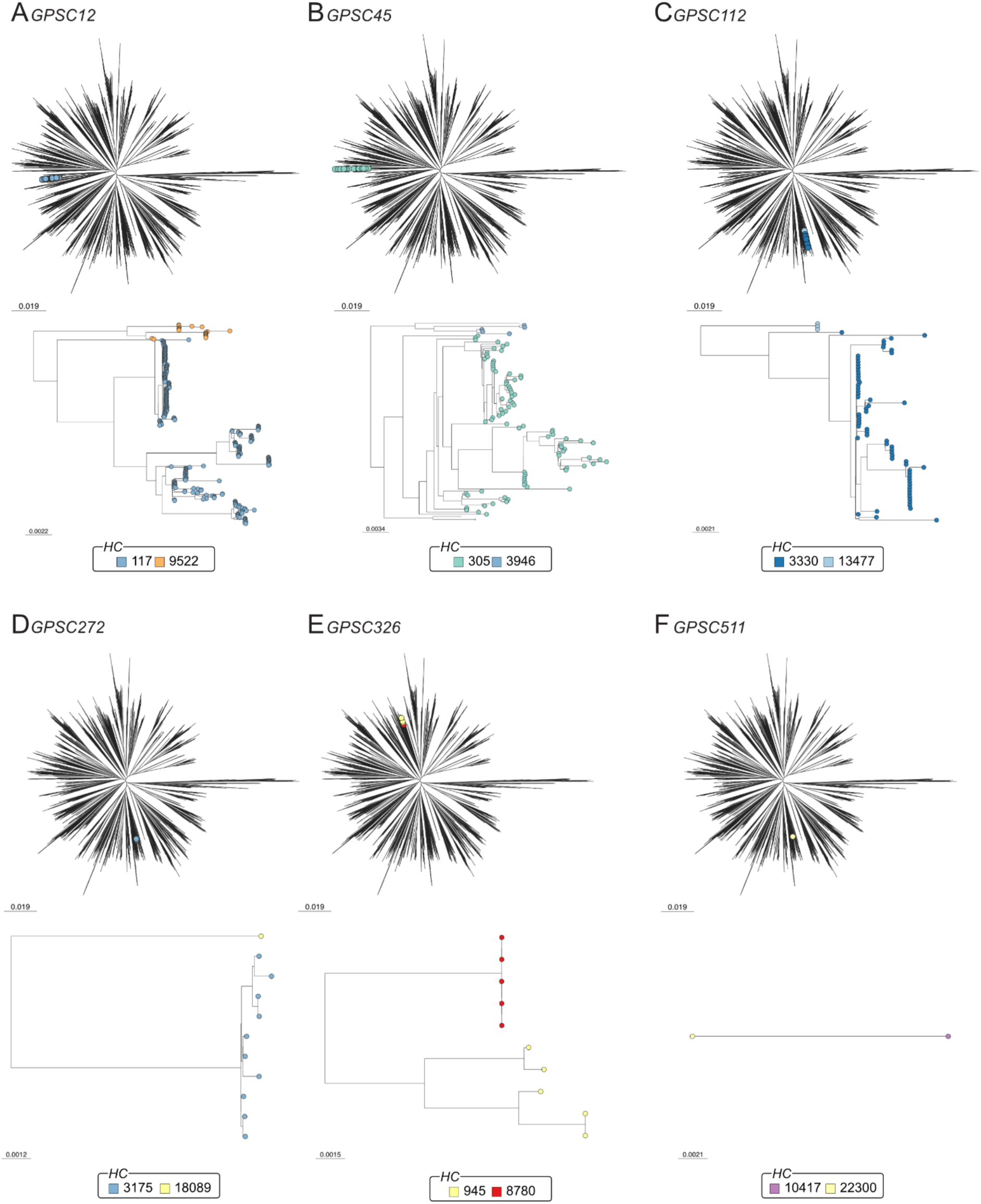
Analysis of the six GPSCs containing two HierCCs. (A) GPSC12, (B) GPSC45, (C) GPSC112, (D) GPSC272, (E) GPSC326, and (F) GPSC511 each contain two HierCCs. The genomes within each GPSC cluster in species-wide phylogenies, and in the subtrees the separation by HierCC was evident.

### HierCCs and LIN Lineage Discordance

Both HierCCs and LIN lineages are based around cgMLST. However, they have different core gene schema and different criteria for defining separate lineages; for HierCCs, clustering is defined as the allelic distance which provides the maximal silhouette score for the dataset [33], whereas LIN lineages are based on the percentage allelic difference between the genomes. In the subset of data that was assigned a LIN lineage from the Pneumococcal Genome Library (PGL) (n = 17,322) [18], there were 474 different HierCCs and 609 different LIN Lineages. There was a mean of 1.00 HierCCs per LIN lineage, and 1.29 LIN lineages per HierCC, showing that LIN lineages are more subdivided than HierCC assignments. As both methods are cgMLST based, we decided to investigate the discrepancies between HierCCs and LIN lineages.

Only one LIN lineage contained multiple HierCCs; this was 0_61_3 (n = 57), which contained members of HierCCs 7 (n = 55) and 510 (n = 2) (**Figure 6A**). All of the genomes clustered together in the species-wide phylogeny, however the cluster is polyphyletic, with closely related genomes being assigned to lineage 0_61_4 (**Figure 6C**). However, HierCC7 and HierCC510 were both part of distinct clades in the subtree (**Figure 6C**). Finally, the GPSC assignments (GPSC94, n = 55; GPSC324, n = 2) were entirely concordant with the HierCC assignments (**Figure 6D**), supporting the division of these genomes into two clusters.

**Figure 6.**
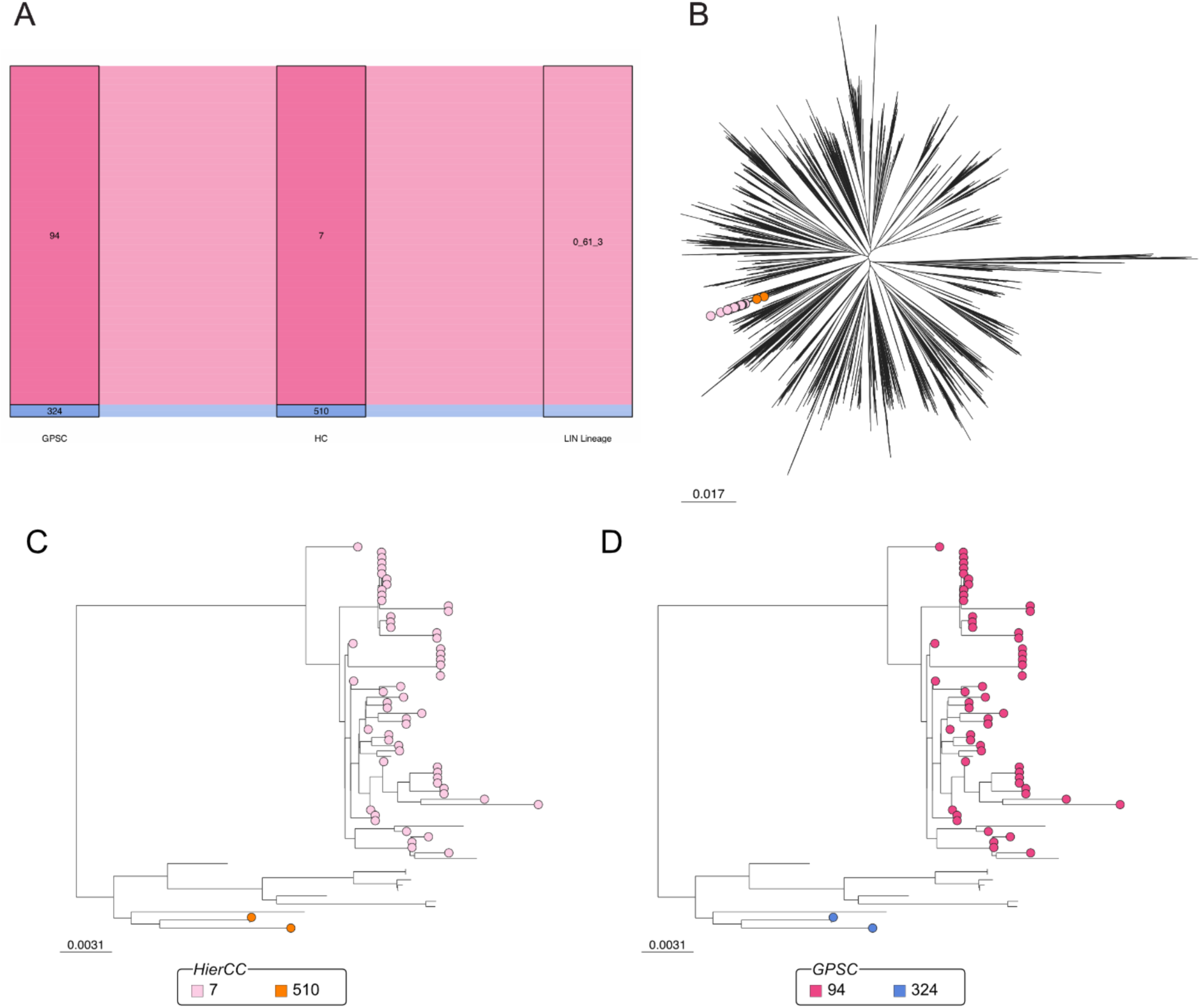
Analysis of genomes assigned to LIN lineage 0_61_3. (A) Both GPSC and HierCC assignments agreed that lineage 0_61_3 should be further divided into two clusters. (B) In a species-wide tree, the genomes were in the same branch. (C) The two HierCCs within 0_61_3 split into two different branches of the subtree. Unlabelled branches represent genomes in LIN lineage 0_61_4. (D) The same is true for the GPSCs. Again, unlabelled branches represent genomes in LIN lineage 0_61_4.

HierCC885 and HierCC390 both contained ten LIN lineages, but as they contained multiple CCs and GPSCs they are discussed earlier in the paper, and the figures can be found in the supplementary.

### CCs Assignment Discordance

As we used data from the PGL to assign LIN barcodes where possible, it is possible to compare the differences between CC assignments in our dataset, and CC assignments from the PGL dataset. There was a 14.5% mismatch between CCs assigned in our dataset and the PGL dataset, showing that the CC nomenclature is contingent on the sample collection. The E-burst analysis showed that large CCs, such as CC156, CC63, and CC185, are formed through connections of smaller clusters by single sequence types (**Supplementary** Figure 15).

## Discussion

Accurate clustering of pathogen genomes is a key part of defining the population structure, as it is likely that closely related isolates may have similar clinical properties such as the disease potential [3], transmissibility [4], and antimicrobial resistance [3]. Additionally, it can also be useful for identifying transmission of *S. pneumoniae* within a population during an epidemic. Additionally, consistent nomenclature is important when communicating results in the global context. Clustering needs to be high resolution, such that differences between groups can be distinguished, but not so fine grained that each cluster only has one or two members, and so the benefits of clustering are lost.

In this study, we used a large and highly diverse global collection of high-quality *S. pneumoniae* genomes to compare the clustering performance of k-mer based PopPUNK in comparison to MLST based clonal complex assignments, and cgMLST based HierCC and LIN lineage assignments. AMI scores between the different clustering methods shows that the results are broadly consistent with each other. However, where they do differ, CCs tend to give the poorest reflection of the population structure based on the species-wide phylogeny, with genomes within single CC could be phylogenetically distant from each other. Additionally, differences in the collection of genomes used to assign CCs may lead to differences in cluster names. Such differences could present a barrier to effective genomic surveillance of *S. pneumoniae*. The two cgMLST methods, HierCC and LIN Lineages, as well as k-mer based PopPUNK, are highly concordant, and their clustering assignments are validated by phylogenetic analysis and pangenome. Therefore, each of these three methods offer a valid approach to defining the population structure of *S. pneumoniae* genomes.

Each of these methods has their own advantages and disadvantages. All cgMLST methods require the definition of a core genome schema, which can differ from study to study depending on the cut-off of prevalence of the gene within the population used to determine a gene as core, and the collection of genomes used to determine the scheme. This can lead to different core genome profiles being used between studies, and potentially different assignments in clustering, as shown by discrepancies between LIN lineages and HierCC assignments. When considering the global spread of *S. pneumoniae*, consistency between studies is key for the accurate interpretation of results and communication between researchers. Additionally, the definition of core genomes can take a significant amount of time. In contrast to this, PopPUNK does not require a core genome schema, leading to greater consistency between studies and faster results, which can be important when describing epidemiology. PopPUNK is the only approach which makes use of the whole genome, including intergenic regions and the accessory genome, instead of just the core genome. However, it still distinguishes between divergence in the core and accessory genomes. This information can be the difference between one cluster or two, as shown with GPSC9 and GPSC235 in HierCC2.

Another key difference between the methods is the ability for clusters to merge over time as new data are added, which is possible for both HierCCs and GPSCs. This can be of benefit, as new genomes support the grouping of the clusters as a better reflection of the population structure. However, this can lead to inconsistencies developing over time as clusters are absorbed into each other, and previous literature is not updated. This is especially true for HierCC assignments, as GPSC assignments avoid this by showing the merging history of a cluster. For example, the assignment GPSC9;904 shows that GPSC904 has merged into GPSC9, and so can avoid this. In contrast to this, the LIN barcoding system does not allow for merging as each genome is provided with a unique barcode. Therefore, the nomenclature will be more stable long-term. Additionally, although not a factor for *S. pneumoniae* due to the equidistant nature of strains, in other species the barcode nature of LIN assignments allows for the user to easily identify the relatedness between two different LIN barcodes based on the nomenclature, whereas the nomenclature of HierCCs and GPSCs does not reflect the hierarchical clustering. For example, it is easy to tell that lineages 0_21_0 and 0_21_1 are closely related, but not how closely related CC63 and CC8346 are. However, at the time of writing this manuscript, it is not possible to assign LIN barcodes data not already within the PGL database.

This work focused on comparing the performance on a single species only, and cannot be translated to other species without further detailed analysis. We used very stringent criteria to define clonal complexes (single-locus variants, only), and altering this could lead again to different clustering results. However, this again reflects the variability in methods. Additionally, further work investigating the validity of minor groupings, such as those with very few genomes, is required to determine the additional value of further subdivision where methods disagree. A final limitation is that whole genome MLST (wgMLST) was not investigated in this study.

As there are now multiple methods available that use genomic data to provide higher resolution clustering than CC assignments, each with different advantages/disadvantages, our findings show that the research community should transition to using these methods over seven-locus MLST. However, to allow for easy comparison between studies and to avoid making previous literature redundant, the reporting of multiple clustering names should be standardised within research. Look-up tables, such as those available on the GPS project website for PMEN, and GPSC interconversion (https://www.pneumogen.net/gps/#/resources#pmen-clones) could be utilised to help with this. This will allow for a greater sense of collaboration within the field, and improve open research practices.

## Supporting information

Metadata

ClusterAssignments

## Acknowledgements

We would like to thank all members of The Global Pneumococcal Sequencing Consortium for their collaborative spirit and determination during the task of sampling, extracting, and sequencing this dataset. We also would like to thank all the clinicians for submitting specimens, and the Wellcome Sanger Institute sequencing facility for sequencing them. Finally, we would like to that the Pathogen Informatics Team at the Wellcome Sanger Institute for technical support of bioinformatic analyses.

---

**Supplementary Figure 1.**
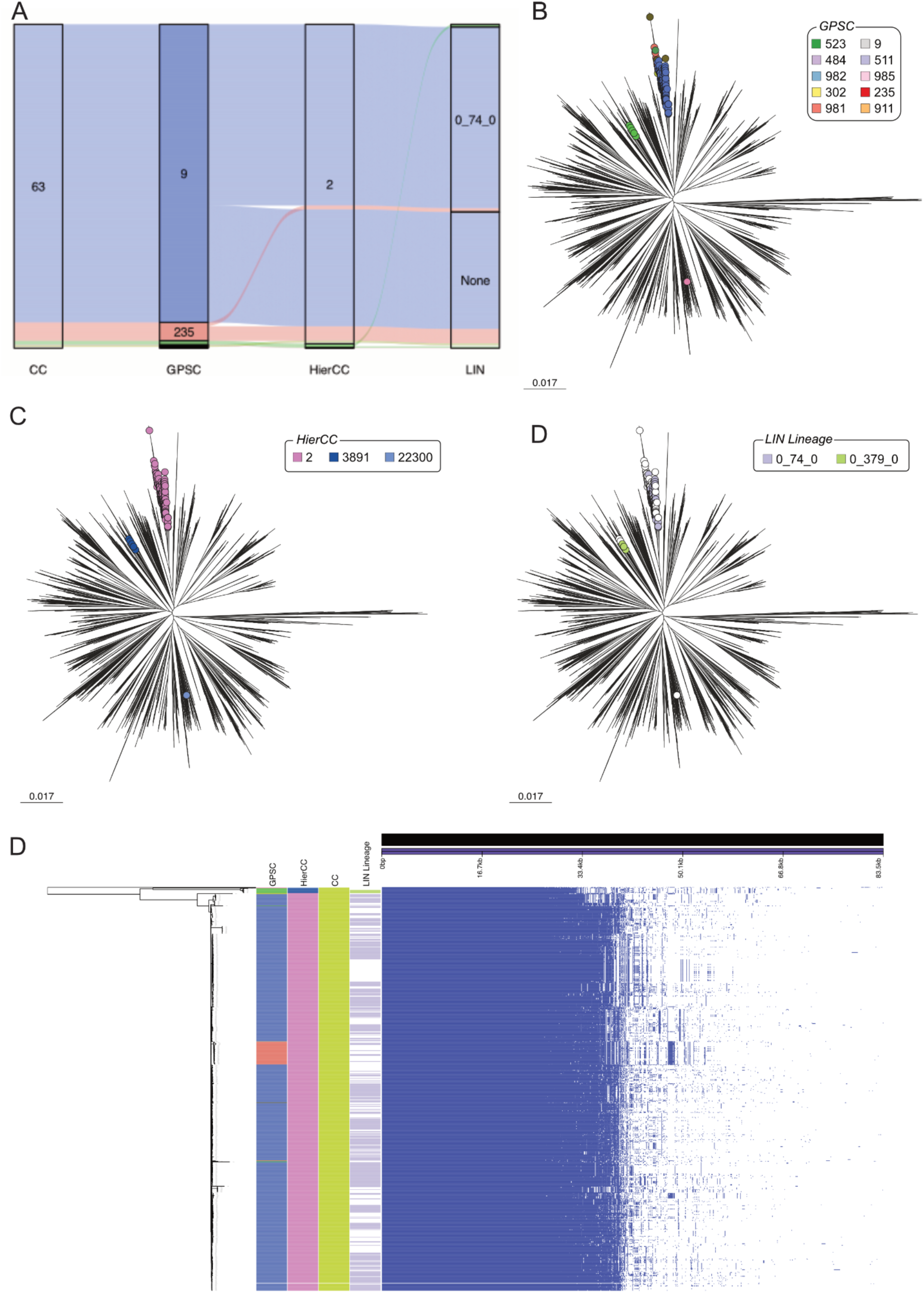
Analysis of genomes assigned to CC63. (A) CC63 is split into multiple GPSCs, HierCCs, and LIN lineages. (B) In a species-wide phylogeny, the genomes clearly belong to distinct branches, reflecting the differences between the genomes and supporting their further subdivision. GPSCs are able to do this. (C) HierCC also clusters the genomes by tree location. (D) The genomes are in different LIN superlineages as well as lineages. (E) Pan-genome analysis supports the separation of CC63 into separate clusters.

**Supplementary Figure 2.**
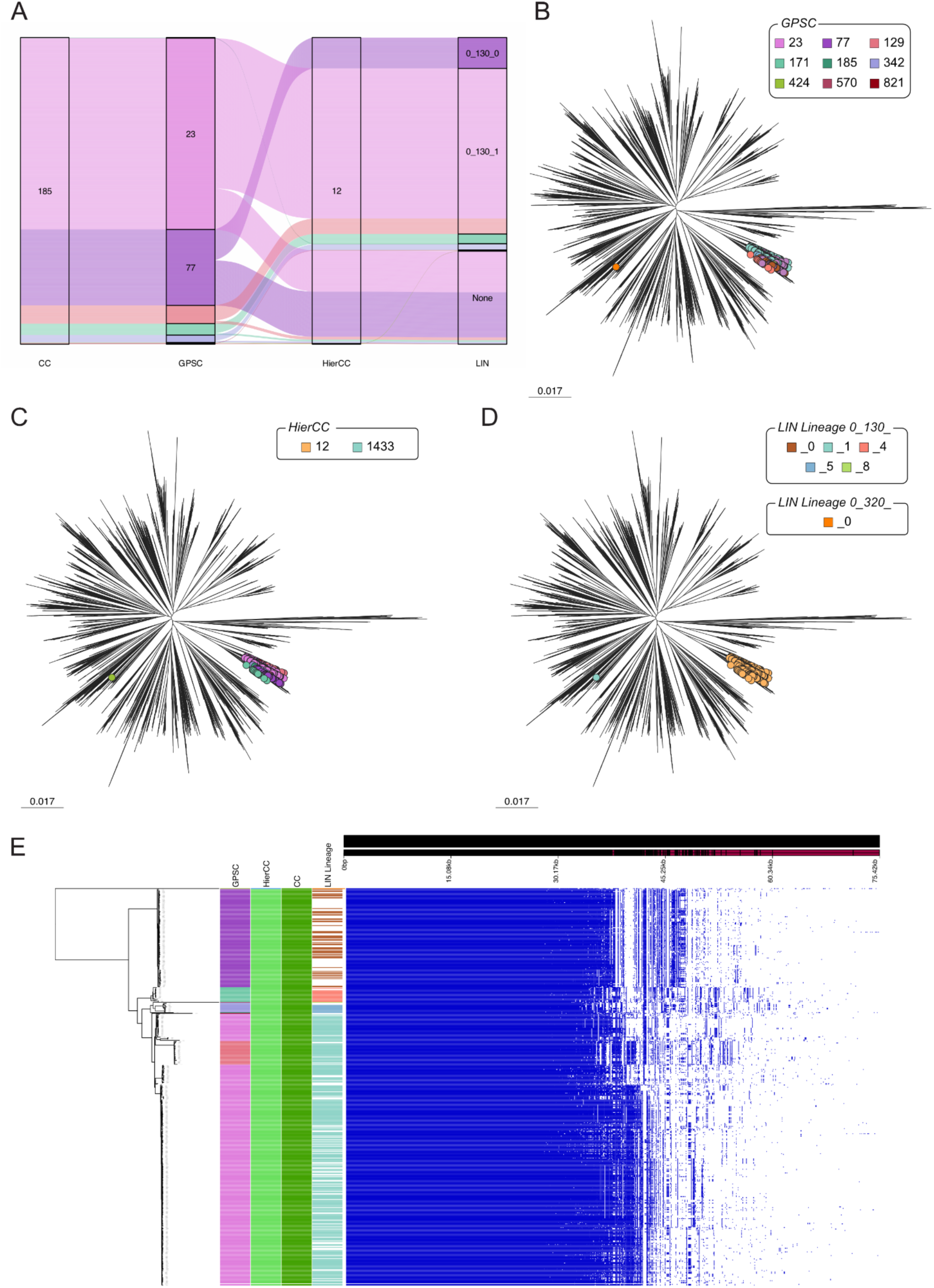
Analysis of genomes assigned to CC185. (A) CC185 is split into multiple GPSCs and LIN lineages, but one HierCC. (B) In a species-wide phylogeny, the genomes clearly belong to distinct branches. GPSCs, (C) HierCC, and (D) LIN lineages are able to distinguish this. The genomes are in different LIN superlineages. (E) Pan-genome analysis supports the separation of CC185 into separate clusters, showing differences in the accessory genomes that have been lost in HierCC assignments, but recognised in LIN lineages and GPSCs.

**Supplementary Figure 3.**
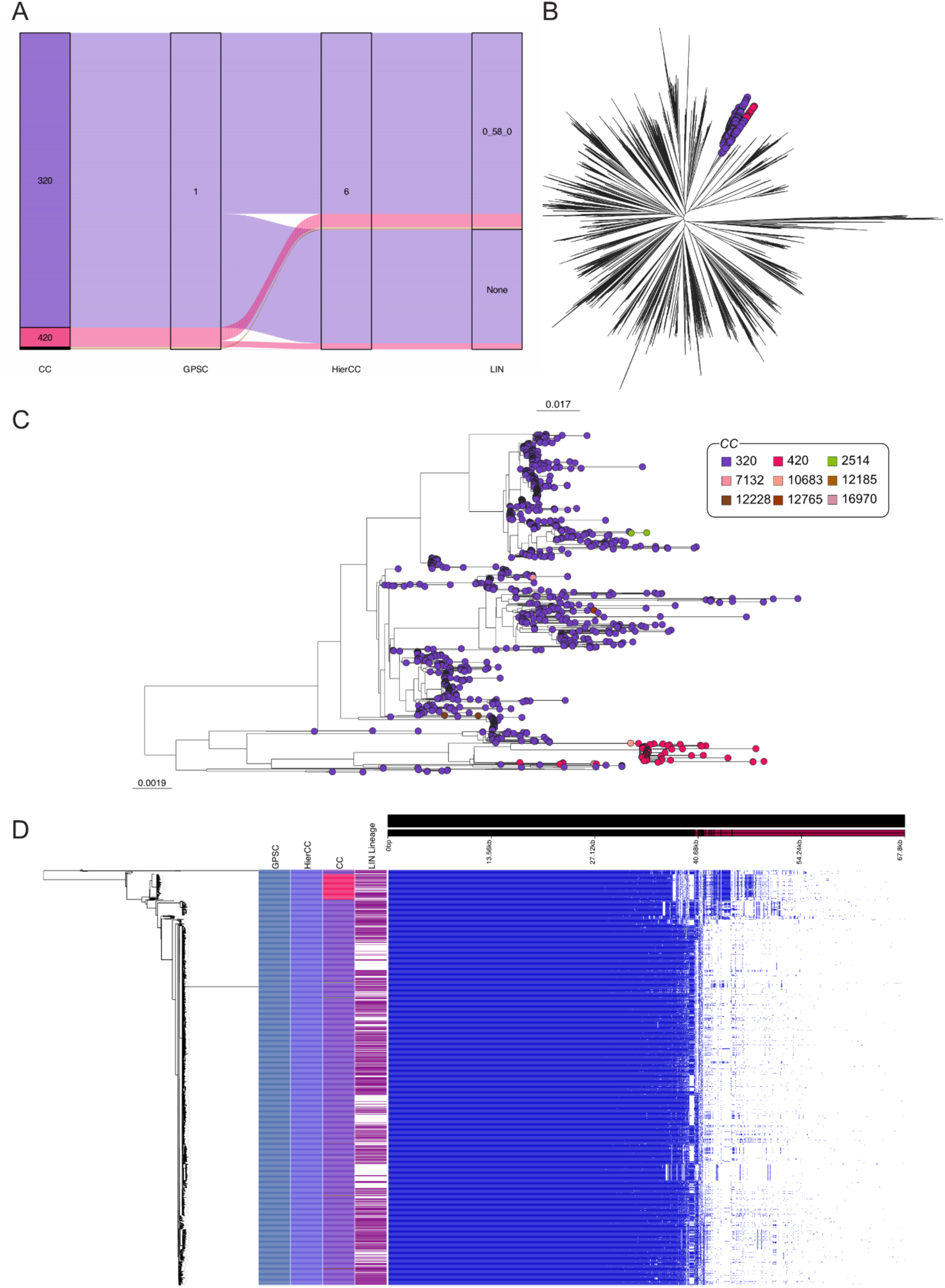
Analysis of genomes assigned to GPSC1. (A) GPSC1 contains multiple CCs, but one LIN lineage and HierCC. (B) In a species-wide phylogeny, the genomes are in the same branch. (C) The subtree coloured by CC shows clustering of two dominant CCs, with many minor CCs scattered throughout. (D) Pan-genome analysis supports the separation of GPSC1 into separate clusters.

**Supplementary Figure 4.**
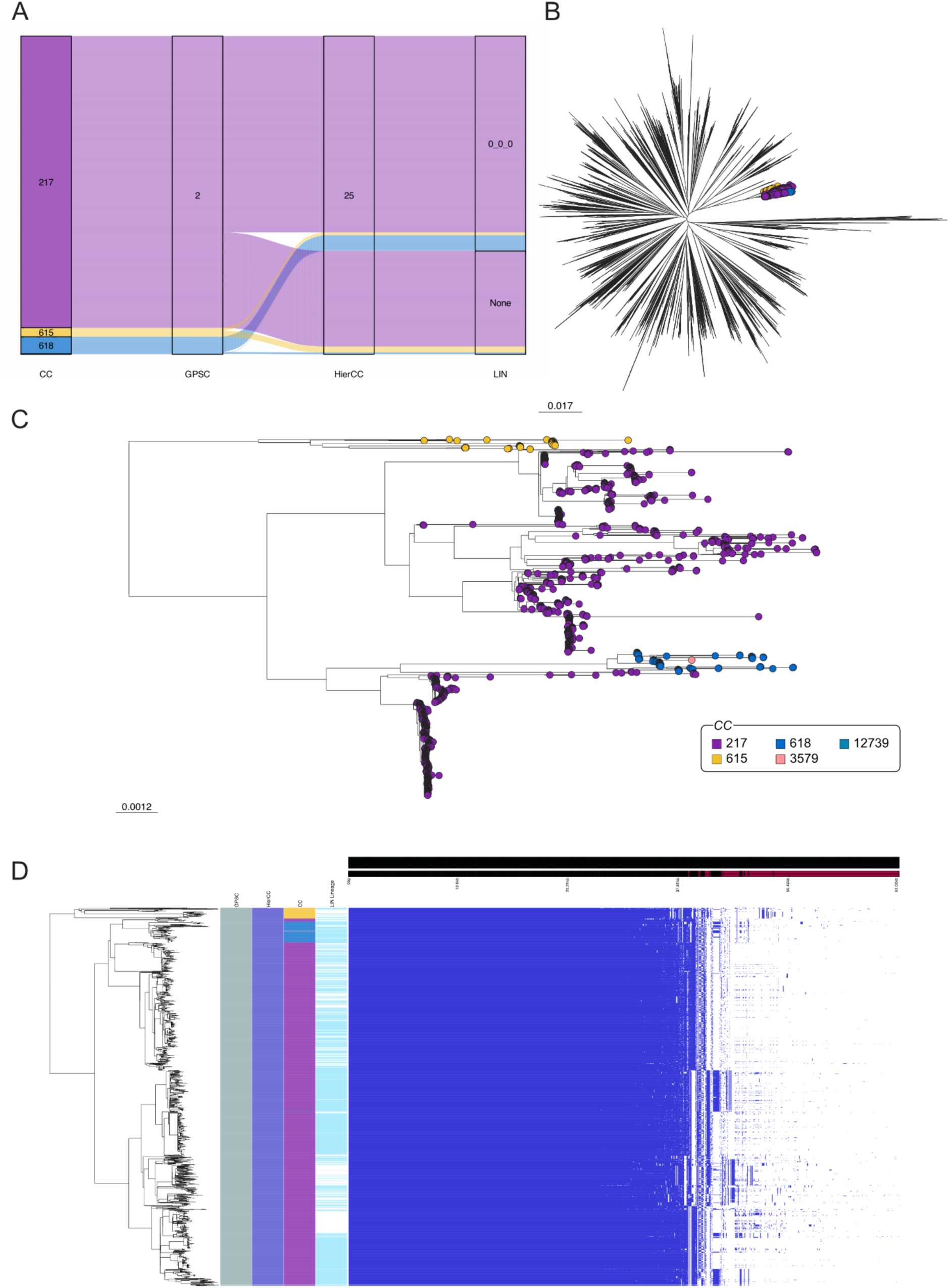
Analysis of genomes assigned to GPSC2. (A) GPSC2 contains multiple CCs, but one LIN lineage and HierCC. (B) In a species-wise phylogeny, the genomes are in the same branch. (C) The subtree coloured by CC shows clustering of the main CCs, with minor CCs scattered throughout. (D) Pan-genome analysis supports the clustering of GPSC1 into a single group.

**Supplementary Figure 5.**
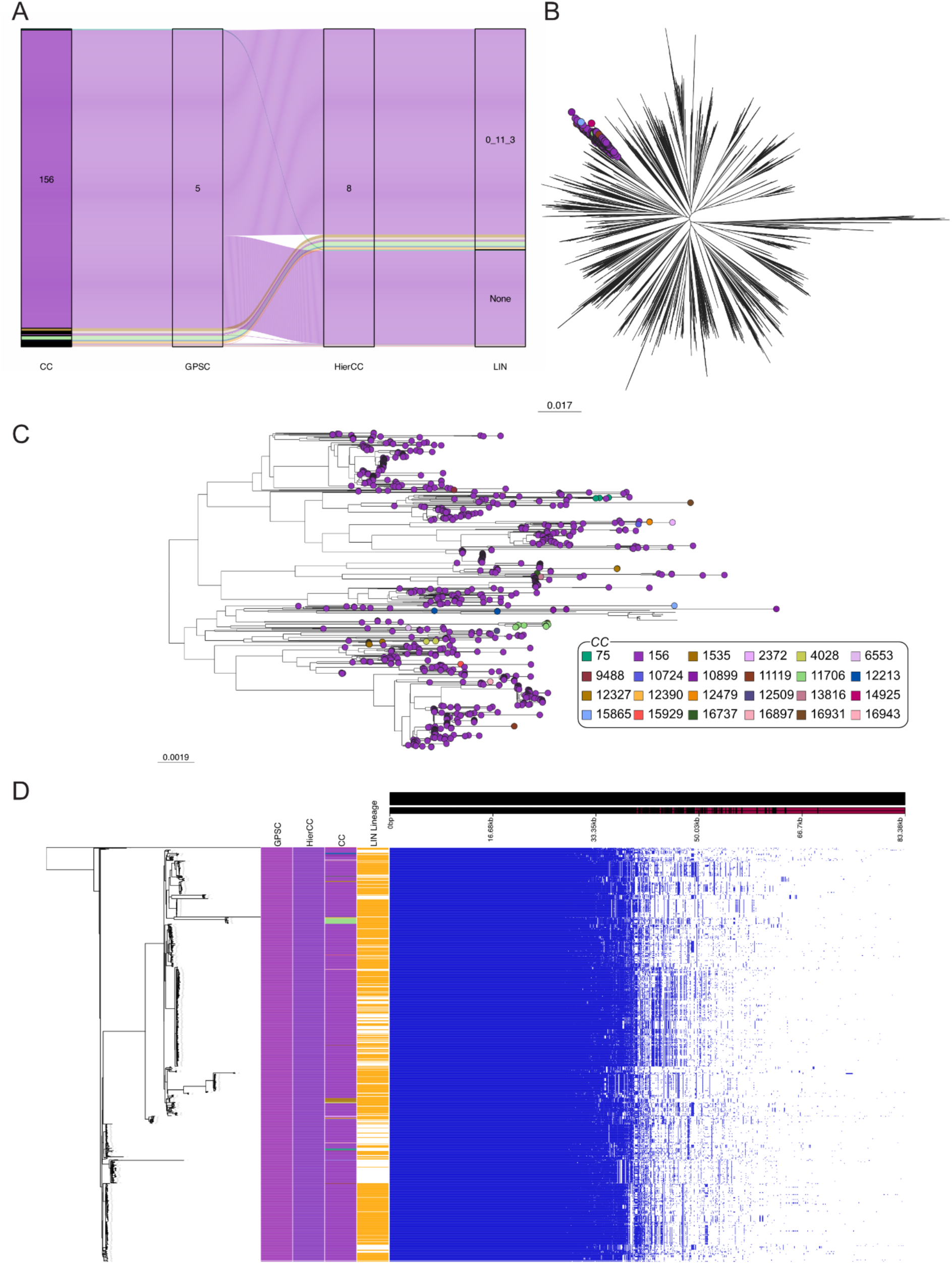
Analysis of genomes assigned to GPSC5. (A) GPSC5 contains multiple CCs, but one LIN lineage and HierCC. (B) In a species-wide phylogeny, the genomes are in the same branch. (C) CC156 is dominant, with many others scattered throughout. (D) Pan-genome analysis supports the clustering of GPSC5 genomes into a single group, and does not provide support for the CC assignments.

**Supplementary Figure 6.**
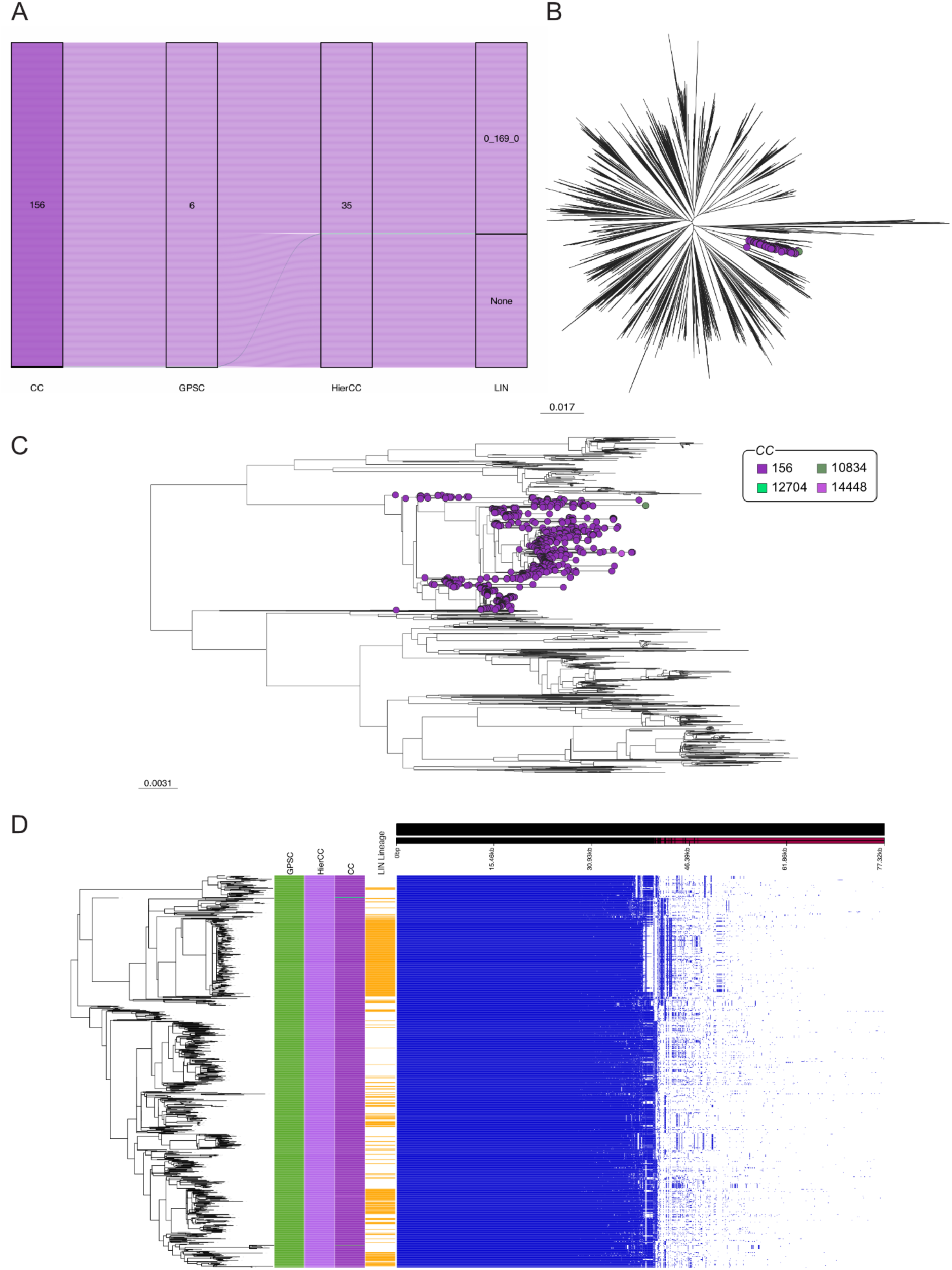
Analysis of genomes assigned to GPSC6. (A) GPSC6 contains a single HierCC and LIN Lineage, but multiple minor CCs alongside the dominant CC156. (B) In a species-wide phylogeny, the genomes are in the same branch. (C) CC156 is dominant, with the other CCs spread across the tree. (D) Pan-genome analysis supports the clustering of GPSC6 genomes into one group.

**Supplementary Figure 7.**
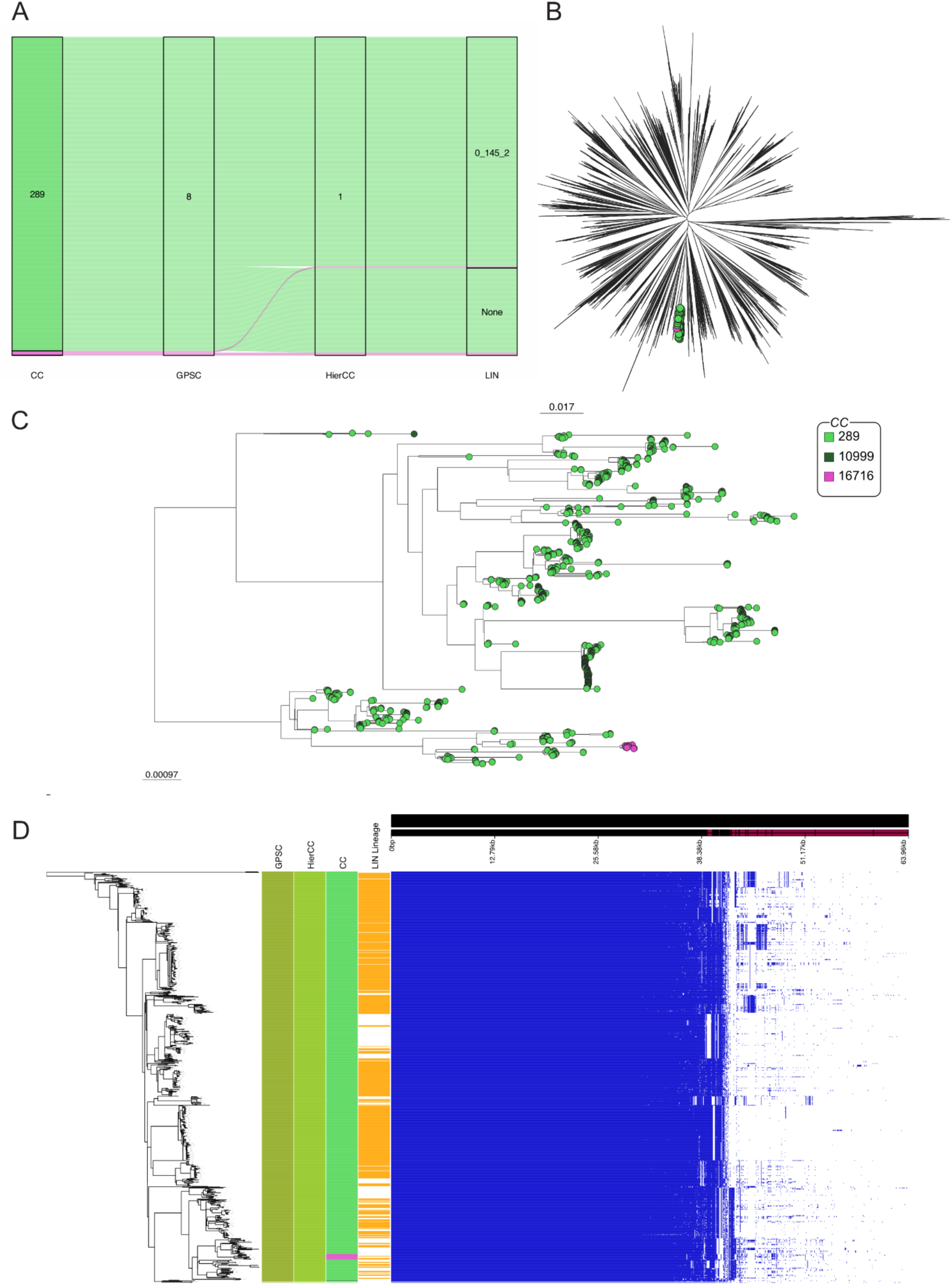
Analysis of genomes assigned to GPSC8. (A) GPSC8 contains a single HierCC and LIN Lineage, but multiple CCs. (B) In a species-wide phylogeny, the genomes are in the same branch. (C) CC289 is dominant, with a small cluster of CC16716 genomes, and a single CC10999 genome. (D) Pan-genome analysis supports the clustering of GPSC6 genomes into one group, and does not support the multiple CCs.

**Supplementary Figure 8.**
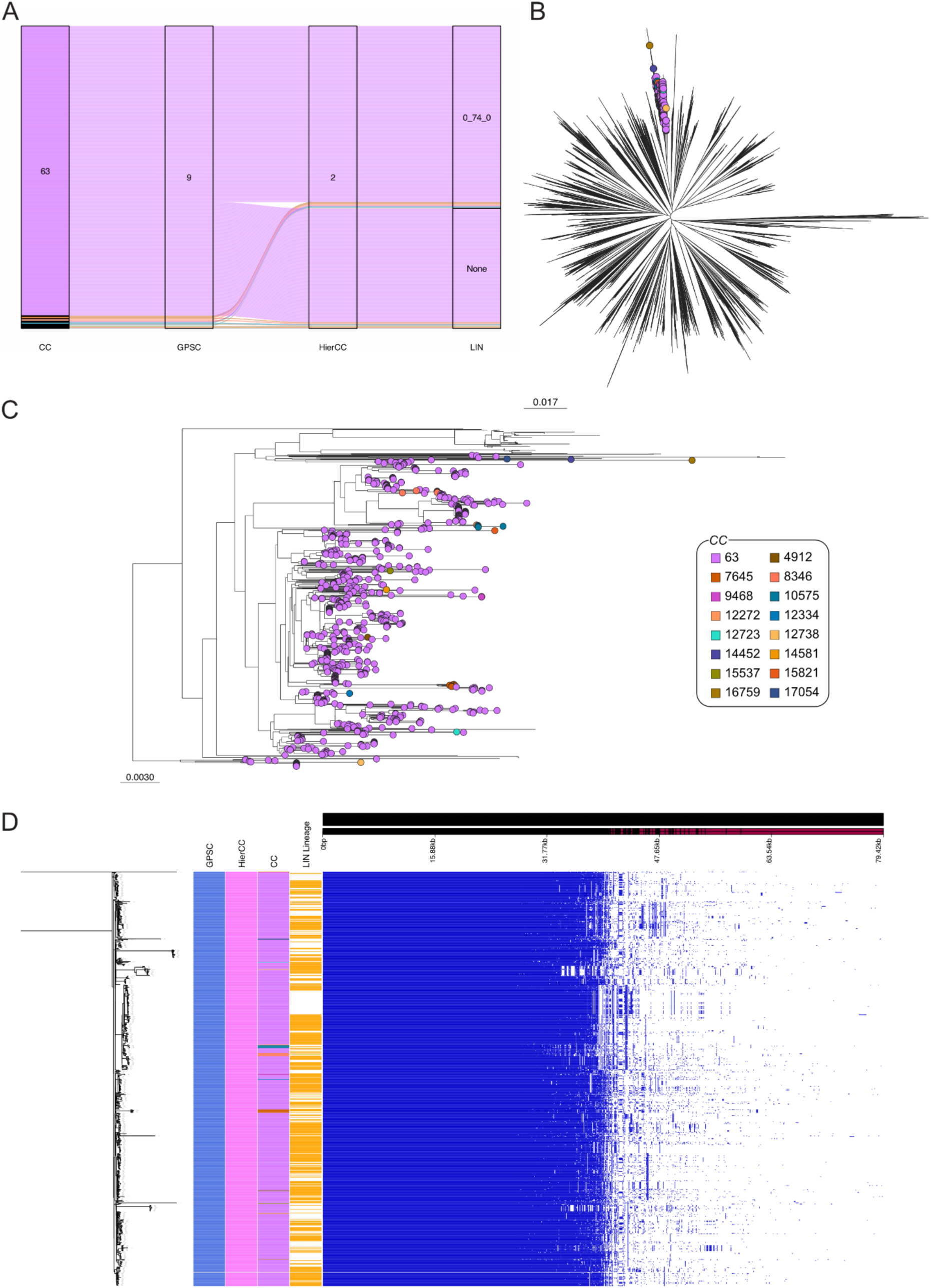
Analysis of genomes assigned to GPSC9. (A) GPSC9 contains a single HierCC and LIN Lineage, but multiple CCs. (B) In a species-wide phylogeny, the genomes are in the same branch. (C) CC63 is dominant, but the phylogeny is very noisy with many minor CCs. (D) Pan-genome analysis supports the clustering of GPSC9 genomes into one group, and does not support the multiple CCs.

**Supplementary Figure 9.**
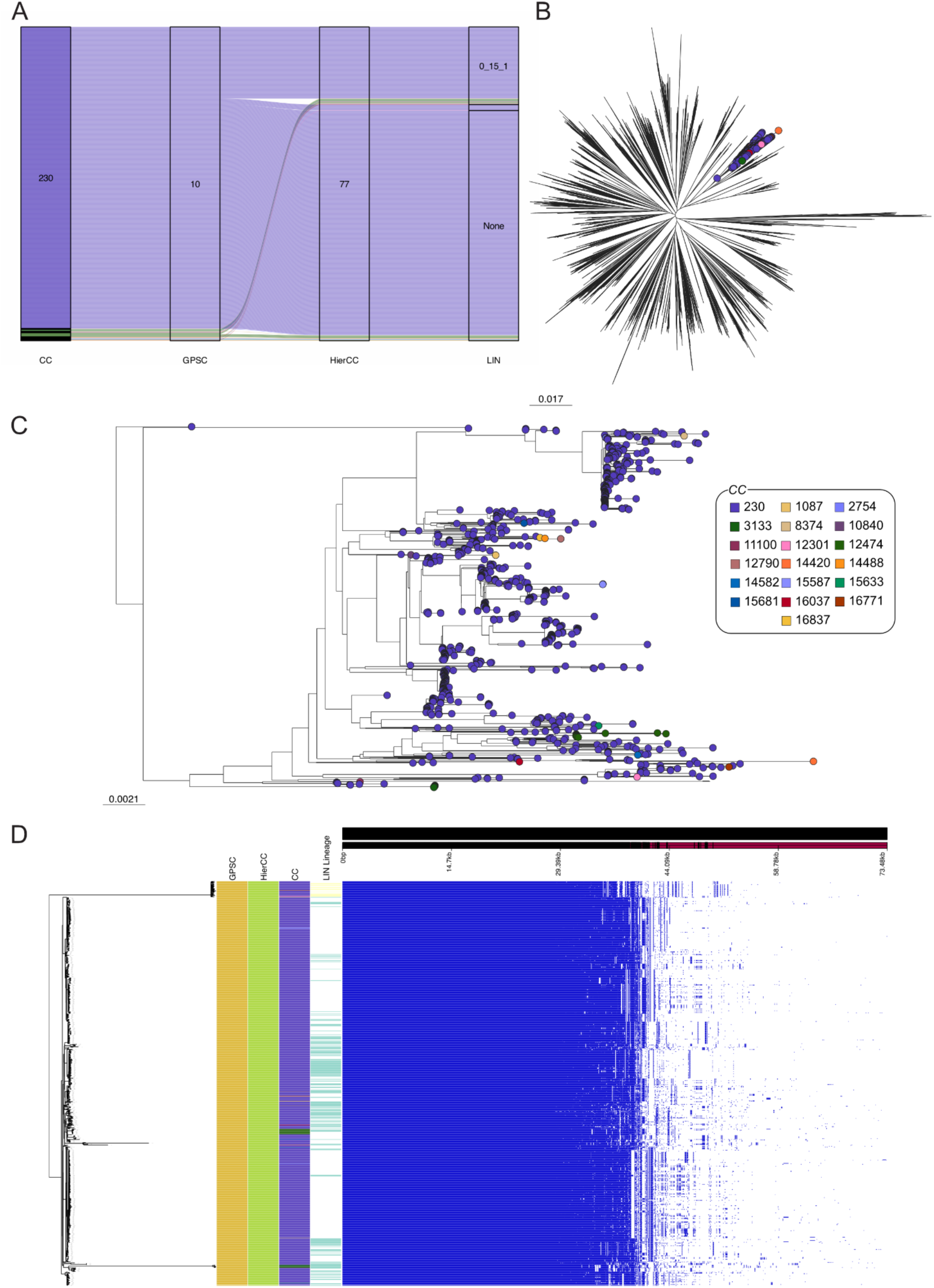
Analysis of genomes assigned to GPSC10. (A) GPSC10 contains a single HierCC but multiple LIN lineages and CCs. (B) In a species-wide phylogeny, the genomes are in the same branch. (C) CC230 is dominant, but other CCs are spread throughout (D) Pan-genome analysis supports the clustering of GPSC10 genomes into one group, and does not support the multiple CCs. However, it does lend support for two the two identified LIN lineages.

**Supplementary Figure 10.**
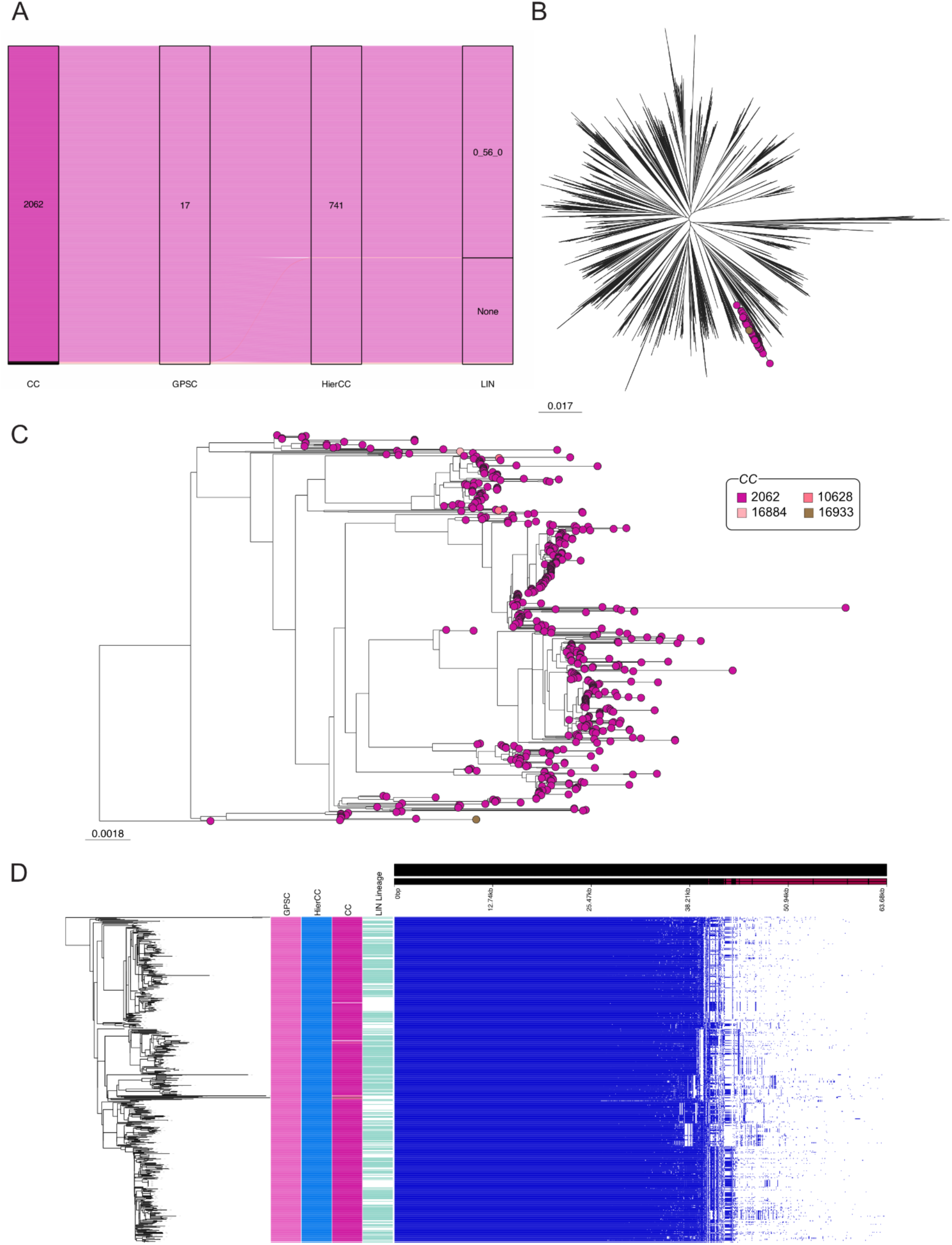
Analysis of genomes assigned to GPSC17. (A) GPSC17 contains a single HierCC and LIN lineage, with four CCs. (B) In a species-wide phylogeny, the genomes are in the same branch. (C) CC2062 is dominant, with the other three represented by very few genomes. (D) Pan-genome analysis supports the clustering of GPSC17 genomes into one group, and does not support the multiple CCs.

**Supplementary Figure 11.**
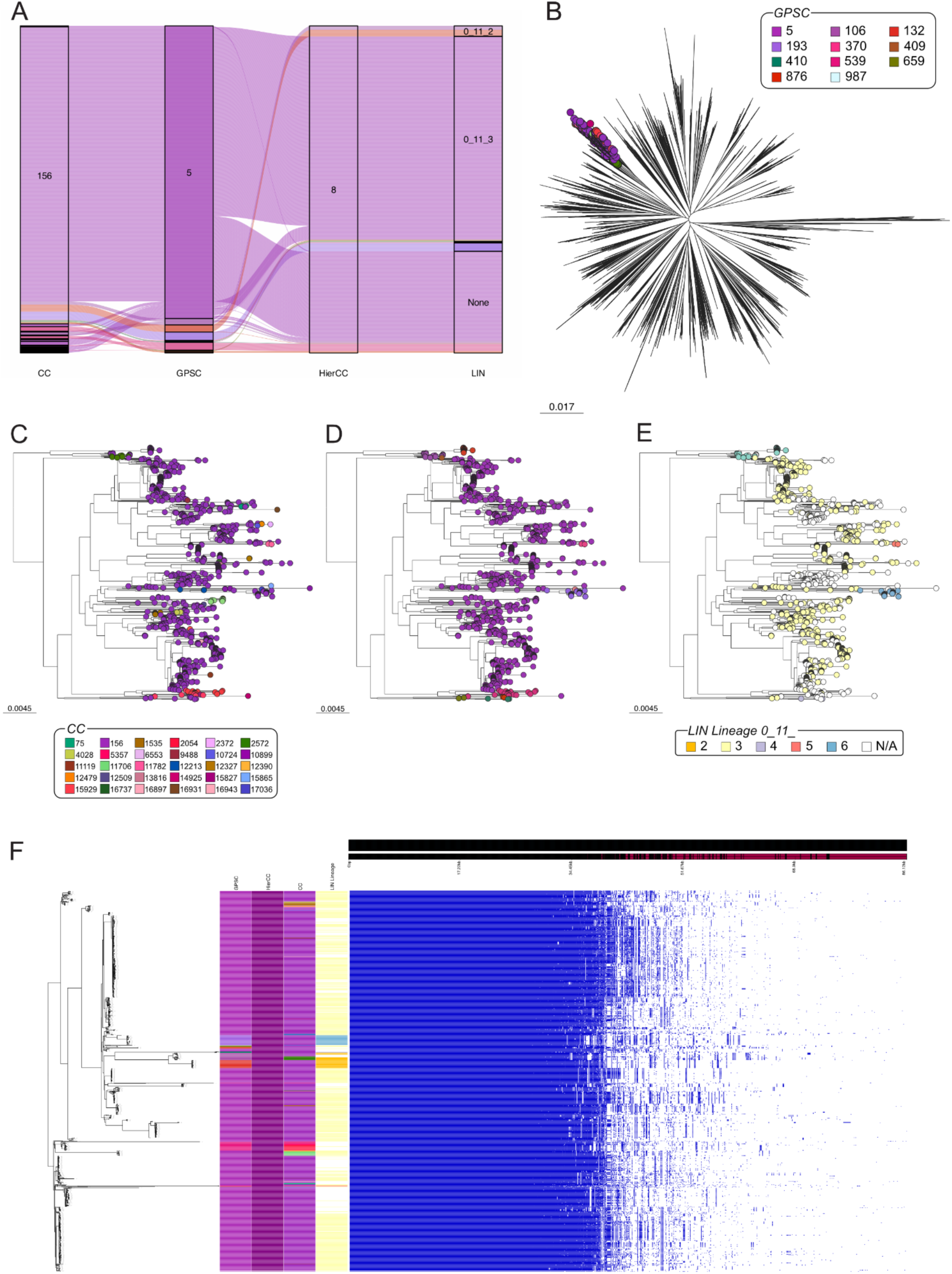
Analysis of genomes assigned to HC8. (A) HC8 contains many GPSCs, CCs, and LIN lineages (B) In a species-wide phylogeny, the genomes are in the same branch. (C) CC156 is dominant, but other CCs are spread throughout and form small clusters (D) There is significant clustering based on GPSC and (E) LIN lineages. (F) Pan-genome analysis does not show huge genomic differences between genomes, however the branch lengths suggest that the division into separate clusters by GPSC assignments is a real effect.

**Supplementary Figure 12.**
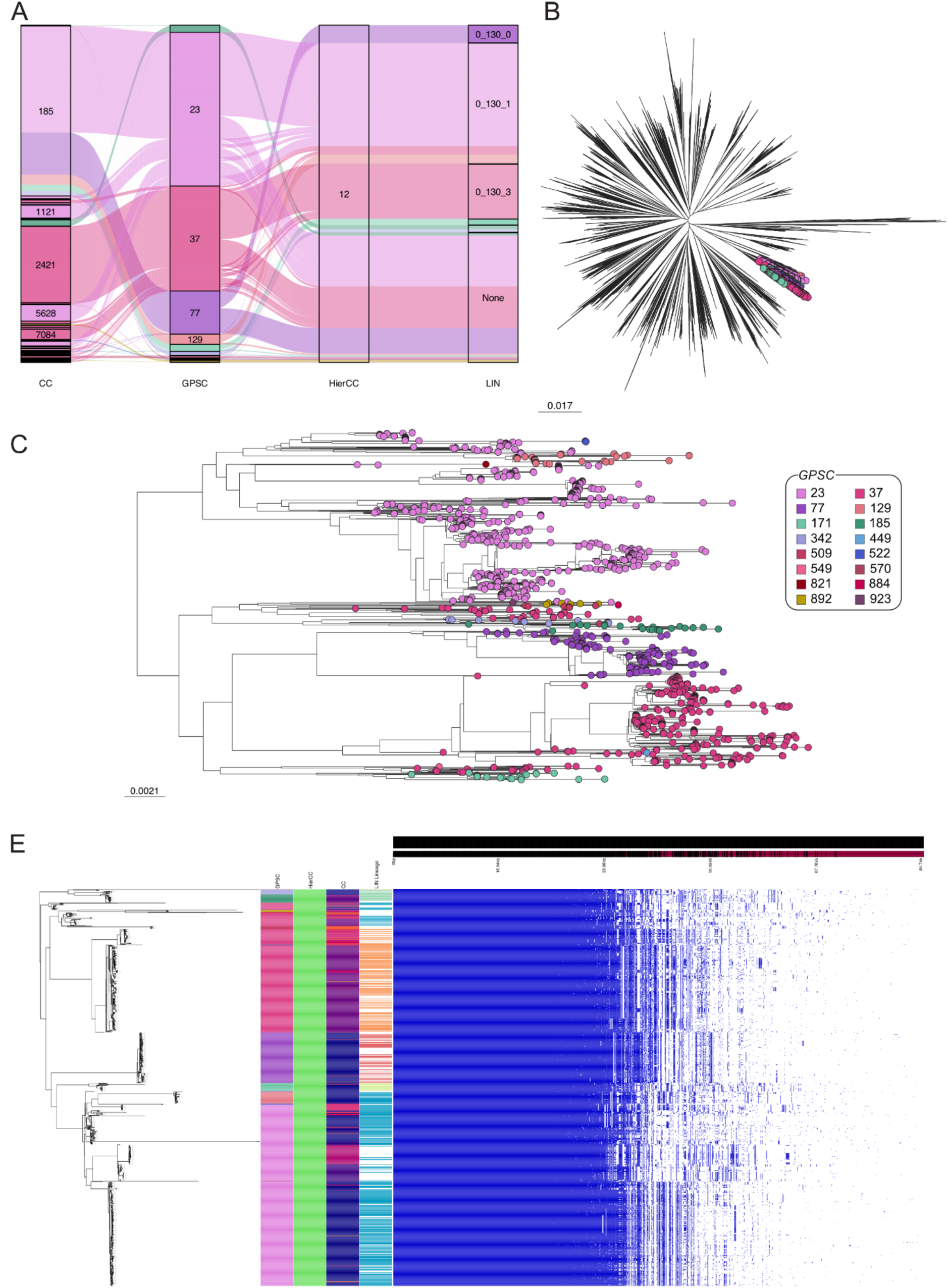
Analysis of genomes assigned to HC12. (A) HC12 contains a huge number of GPSCs, CC, and LIN lineages. (B) In a species-wide phylogeny, the genomes are in the same branch. (C) Clear clustering is seen in the subtree based on GPSC. (D) Pan-genome analysis broadly supports the GPSC assignments, though there are still limited differences between the genomes.

**Supplementary Figure 13.**
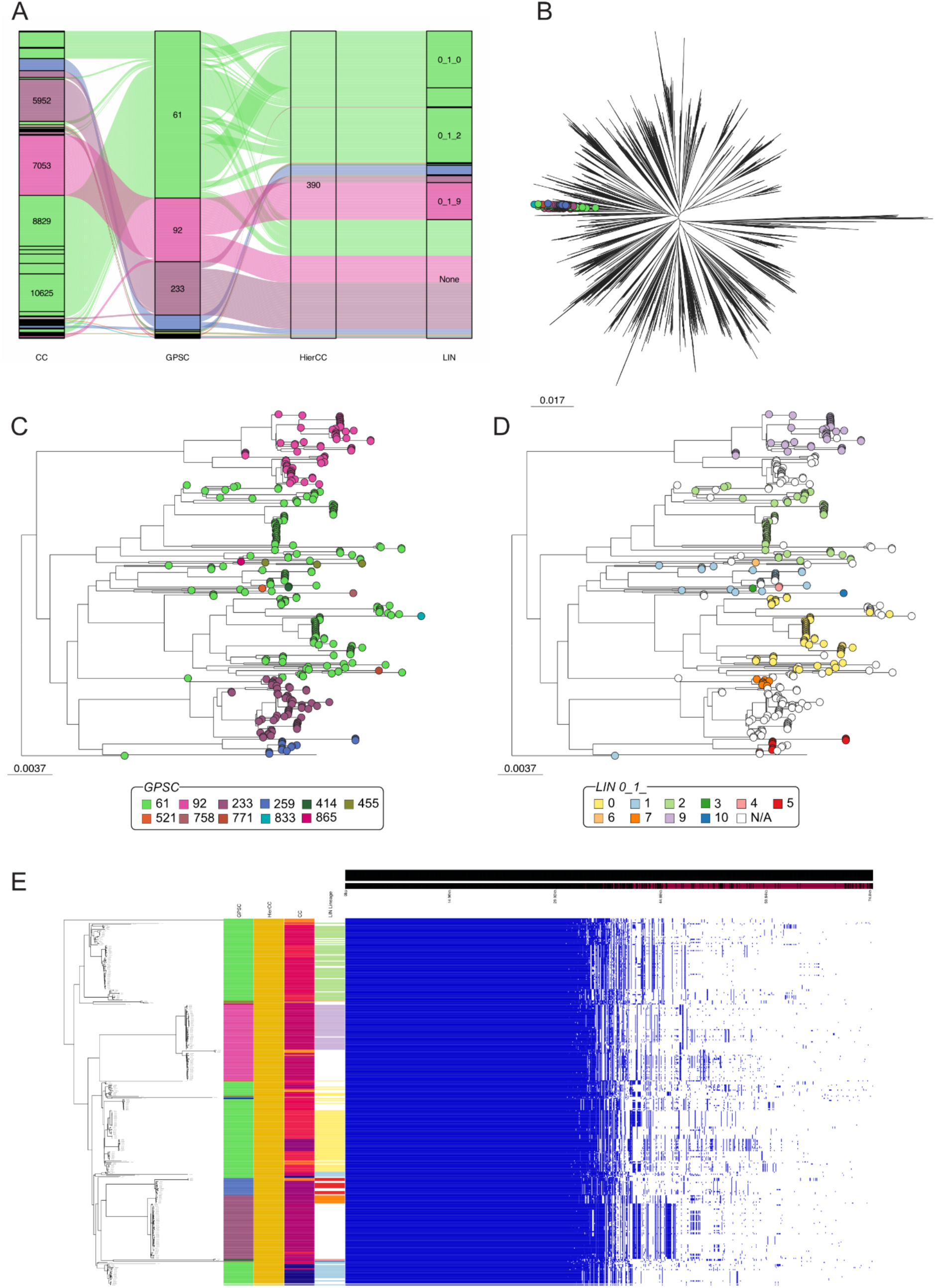
Analysis of genomes assigned to HC390. (A) HC390 contains a huge number of GPSCs, CC, and LIN lineages. (B) In a species-wide phylogeny, the genomes are in the same branch. (C) Clear clustering is seen in the subtree based on GPSC and (D) LIN lineage. (E) Pan-genome analysis supports the GPSC assignments.

**Supplementary Figure 14.**
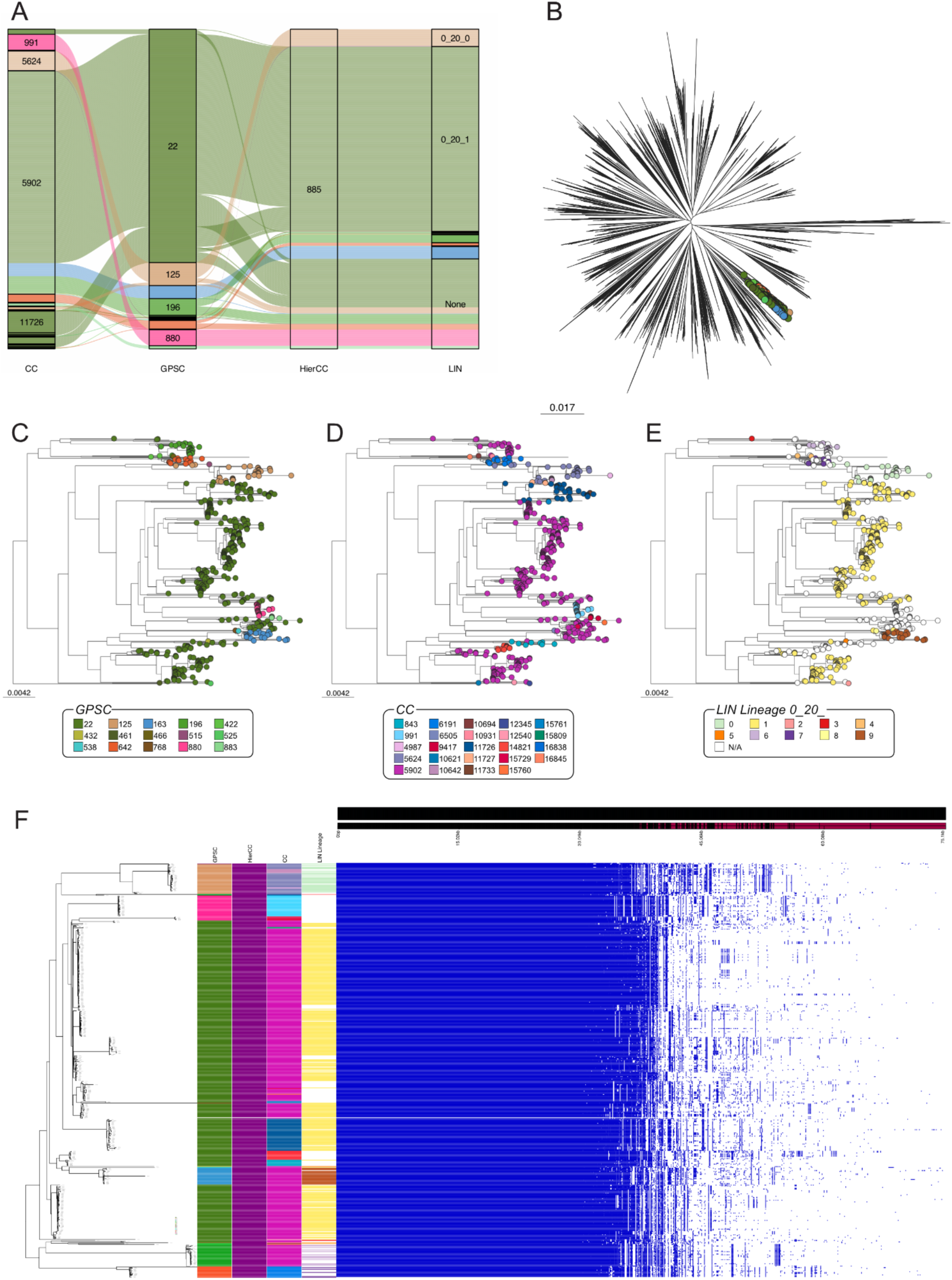
Analysis of genomes assigned to HC885. (A) HC885 contains many GPSCs, CC, and LIN lineages. (B) In a species-wide phylogeny, the genomes are in the same branch. (C) Clear clustering is seen in the subtree based on GPSC, whereas (D) CC, and (E) LIN lineages do not cluster quite as well. (F) Pan-genome analysis supports the GPSC assignments, as seen by differences in the gene content.

**Supplementary Figure 15.**
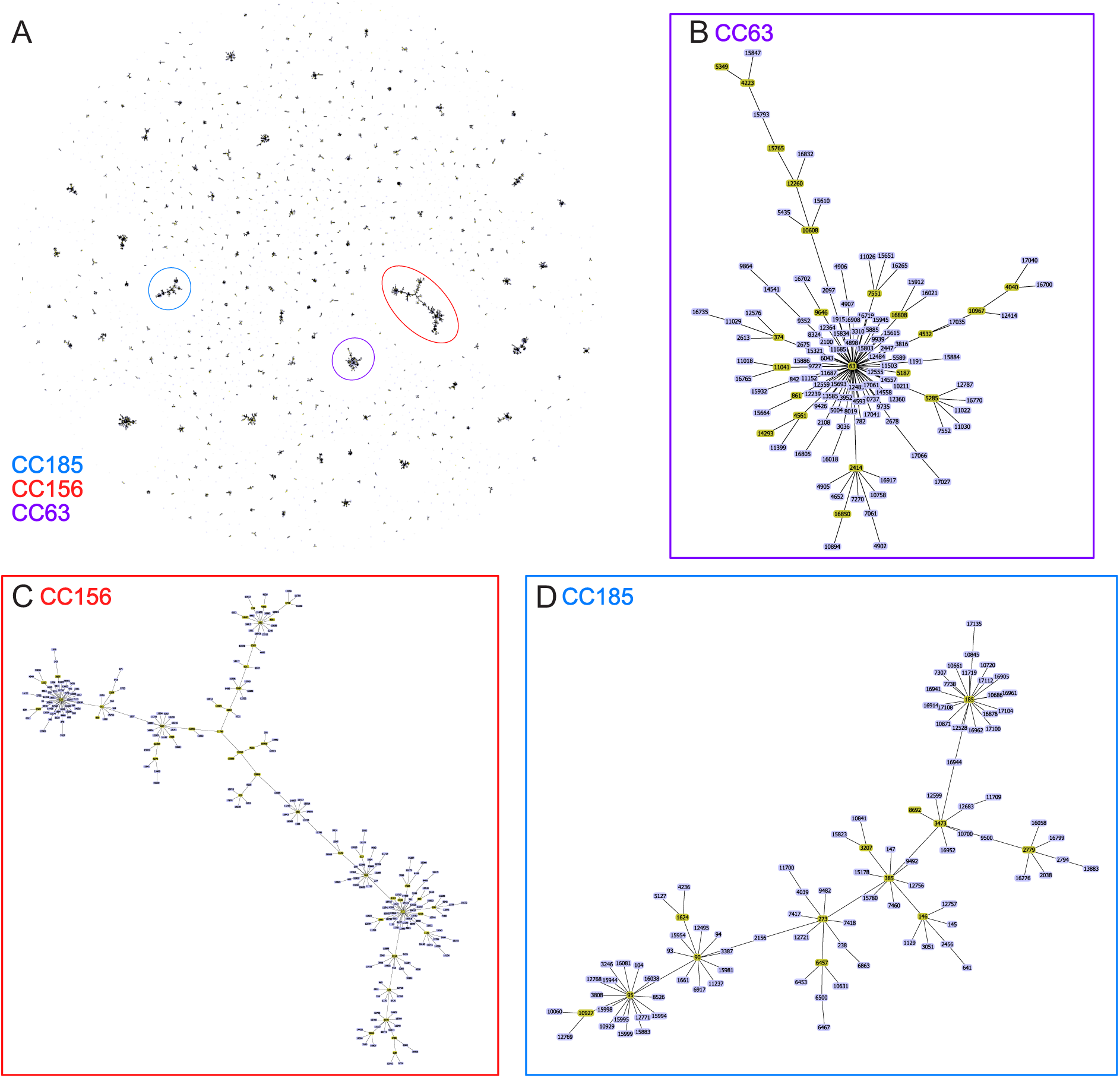
Phyloviz E-burst output for all genomes within the dataset, with a single locus variant cut-off. Each connected cluster represents a CC. (A) A huge number of CCs are seen, with many being singletons or very small. However, larger CCs such as CC63 (B), CC156 (C), CC185 (D) can also be seen. In these larger networks, often a single ST can connect two larger groups of STs.

**Supplementary table 2.** A selection of genes present in GPSC235, but not in GPSC9, despite both sets of genomes being clustered into HierCC2. Taken from Panaroo annotation.

| *Gene* | *Non-unique*<br>*Gene name* | *Annotation* |
| --- | --- | --- |
| group_829 |  | membrane protein |
| group_766 |  | phage protein |
| group_1725 |  | FIG01114970: hypothetical protein |
| group_1703 |  | transposase;transposaseTransposase domain (DUF772) |
| group_1658 |  | glutamate-cysteine ligase |
| group_1637 |  | arsenate reductase |
| group_1630 |  | site-specific recombinase;site-specific recombinaseRecombinase;site-specific recombinaseResolvase N terminal domain |
| group_1629 |  | hypothetical protein |
| pezT_1 | pezT_1 | zeta toxinUDP-N-acetylglucosamine kinaseUncharacterized protein conserved in bacteriaZeta toxin |
| group_1517 |  | FIG01114020: hypothetical proteinReplication initiator protein A (RepA) N-terminus |
| group_1513 |  | transposase;transposaseTransposase DDE domain;transposaseTransposaseTransposase DDE domain |
| immR~~~immR_2~~~immR_1 | immR;immR_2; immR_1 | Cro/CI family transcriptional regulatorHTH-type transcriptional regulator immRPredicted transcriptional regulatorHelix-turn-helix |
| group_1500 |  | ATPases with chaperone activity ATP-binding subunit |
| group_1490 |  | Abortive infection protein AbiGIINucleotidyl transferase of unknown function (DUF1814) |
| group_1480 |  | membrane protein |
| group_1478 |  | FIG01114516: hypothetical protein |
| group_1429 |  | FIG01116986: hypothetical proteinPrgI family protein |
| group_1408 |  | Tn5252 Orf 10 protein |
| group_1386 |  | Tn5253 hypothetical protein |
| group_1349 |  | FIG01114502: hypothetical protein |
| group_1326 |  | site-specific recombinasemultiple promoter invertaseResolvase N terminal domain |
| group_1312 |  | Tn5252 ORF9Bacterial mobilisation protein (MobC) |
| group_1285 |  | trsE-like proteinType IV secretory pathway VirB4 componentstype-IV secretion system protein TraCAAA-like domain |
| group_1278 |  | hypothetical protein |
| group_782 |  | hypothetical protein |
| hpaIIM_1 | hpaIIM_1 | C-5 cytosine-specific DNA methylaseModification methylase HpaIIDNA cytosine methylaseDNA (cytosine-5-)-methyltransferaseC-5 cytosine-specific DNA methylase |
| pezA | pezA | DNA-binding helix-turn-helix proteinAntitoxin PezAhypothetical proteinPredicted transcriptional regulatorputative zinc finger/helix-turn-helix protein YgiT familyHelix-turn-helix domain |
| group_692 |  | hypothetical protein;Uncharacterized protein conserved in bacteria (DUF2326) |
| group_634 |  | site-specific recombinaseRecombinase;site-specific recombinase |
| group_528 |  | DNA primase (type)DNA primase (bacterial type)DNA primase;FIG01116415: hypothetical proteinDNA primase;DNA primase (type) |
| group_478 |  | SNF2 family proteinDNA methylaseDNA phosphorothioation system restriction enzymeSNF2 family N-terminal domain;SNF2 family proteinDNA methylase |
| group_410 |  | Tn5253 hypothetical protein |
| group_374 |  | relaxaseRelaxase/Mobilisation nuclease domain |
| group_348 |  | FIG01114872: hypothetical protein |
| ssp5 | ssp5 | Agglutinin receptorSSP-5Predicted outer membrane proteinadhesin isopeptide-forming adherence domainGlucan-binding protein C;Agglutinin receptorSSP-5adhesin isopeptide-forming adherence domainGlucan-binding protein C |

